# Decoding murine corneal epithelial specification and homeostasis by single-cell spatial transcriptomics with scRNA-seq enrichment

**DOI:** 10.64898/2026.05.08.723186

**Authors:** Dina Javidjam, Meri Vattulainen, Neil Lagali, Petros Moustardas

## Abstract

Investigation of gene regulatory programs underlying corneal epithelial cell specification and homeostasis is essential for understanding how the cornea maintains vision. Here, we describe the use of true single-cell resolution spatial transcriptomics (ST), enriched with full-tissue single-cell RNAseq (SC), to improve spatial resolution and enhance cell cluster size up to 65-fold and per-cell transcriptomic depth up to 17-fold. This enabled cell type specification across the full differentiation trajectory from limbal stem cells (LSC) to superficial corneal epithelium and identification of an activated signature (*Atf3, Zfp36, Gsta4 and Dapl1*) marking differentiation-primed states across multiple cell types, including a major activated intermediate epithelium (AIE) population. Validation using ST data from murine corneas at different postnatal ages and multiple human SC datasets confirms a large AIE population, which spatial localization and transcriptomic profiling suggest is an active intermediate state distinct from quiescent wing cells. Sub-clustering further revealed early (*Sox9, Hes1*), proliferative (*Mki67, Top2a*) and mature (*Ccdn1, Dapl1*) transient amplifying cell subpopulations and four LSC subpopulations, including putative active (*Atf3, Socs3, Zfp36*), quiescent (*Gpha2, Ifitm3, Cd63*) and *Apoe*-specific. Direct ST-to-SC comparison revealed enhanced axonal processes and genes (*Sema3f, Sema 4d, Pax6*) and cell-cell adhesion and cell-matrix markers (*Itgb4, Tns4, Tjp3*) in ST data, suggesting cell dissociation from tissue in SC masks epithelial innervation, adhesion and barrier functions. Our findings identify and localize key transcriptional programs *in situ*, prompting a re-evaluation of epithelial states in scRNA-seq data.

## Introduction

The corneal epithelium forms the outermost layer of the transparent cornea, whose renewal is sustained by limbal stem cells (LSCs) located at the corneoscleral limbus^1^. LSCs give rise to progenitor, activated and differentiated cell populations that comprise the stratified epithelium and confer its critical barrier and transparency functions, and importantly, an ability to effectively regenerate following injury, to preserve proper vision. The maintenance of corneal homeostasis relies on tightly controlled regulatory mechanisms, making detailed insight into cellular specification and spatial organization essential for better understanding corneal health and disease.

Substantial efforts have been directed toward uncovering the complexities of LSC heterogeneity and differentiation dynamics, with implications for corneal wound healing and transparency maintenance across a range of diseases. Transcriptomic profiling methods such as high-throughput single-cell RNA sequencing (scRNA-seq, hereafter SC) have revealed the complexity of corneal epithelial subpopulations and the fine-tuned hierarchical differences particularly amongst stem and progenitor populations in mice^2–7^ and in humans^8–12^.

SC studies, however, are influenced by cell dissociation-induced artifacts^13,14^ and loss of spatial information, which can distort native transcriptional profiles and reduce applicability to the *in vivo* context, ultimately limiting our understanding of transcriptional programs of cell specification, intercellular and cell-matrix interactions, and regional differences within the cornea. Alternatively, high-resolution spatial transcriptomic (ST) technologies preserve tissue architecture and spatial context of cells within their native environment. Increased spatial resolution in thin tissue sections is nonetheless typically accompanied by reduced gene detection capacity, highlighting an inherent trade-off compared to SC^15^. To address these limitations, integrative approaches exploiting the strengths of both ST and SC are required to enable spatial mapping of defined cell populations while providing insight into how these populations interact to support tissue homeostasis. Current computational integration strategies, however, remain suboptimal^16^, as no current ST platform simultaneously achieves true single-cell-resolution with the depth and comprehensive transcriptome coverage of SC.

Here, we sought to elucidate the deep transcriptomic and spatial specification of the postnatal mouse corneal epithelium in homeostasis by leveraging the strengths of both ST and SC. To achieve this, we integrated SC with high-resolution ST using a Single-Cell Variational Inference (scVI)–based framework^17^, coupled with a custom workflow enabling true single-cell segmentation of ST data. Our approach enabled the deconstruction of deep transcriptomic states comprising the full murine corneal epithelium, while retaining high-resolution spatial information. This leads to a robust identification of cell populations and an active cell signature within the corneal epithelium suggesting re-evaluation of prior SC data. The approach also enables smaller cellular subpopulations with distinct transcriptional profiles to be identified for deeper investigation, such as functionally distinct stem cell subtypes. We also highlight advantages of fresh ST tissue samples in preserving contextual information such as cell-to-cell adhesion, barrier, and junctional programs, as well as cell-to-matrix and corneal innervation features undetectable in SC data. Finally, we show how our detected cell states and spatial information could be extended to new ST samples and postnatal ages outside the original integration dataset, highlighting broader applicability of the robust cell state classifications.

## Materials and Methods

### Ethics statement

This study was conducted after receiving approvals from the Linköping Regional Animal Ethics Review Board, Linköping, Sweden (Protocol numbers 10940-2021 and 17409-2023) and was carried out in accordance with the ARVO statement for the Use of Animals in Ophthalmic and Vision Research as well as the EU Directive 2010/63/EU on the protection of animals used for scientific purposes.

### Animals

Wild type mice on a 129S1/SvImJ background (MMRRC stock #050624-MU) were housed at Linköping University under standardized conditions (23 °C, 40–60% relative humidity, 12-hour light/dark cycle). Six corneas from three mice aged 4 months were collected for pooled SC. Additionally, a total of four corneas were collected for ST at two months or four months of age (two corneas from two mice per timepoint). Mice were first anesthetized with intraperitoneal ketamine-xylazine injection (65 mg/kg ketamine, 10 mg/kg xylazine diluted in PBS for 10 μL/g of weight injection) and thereafter euthanized through cervical dislocation.

### Single-cell RNA sequencing (SC)

Mouse corneas were freshly dissected while immersed in cold PBS from enucleated eyes and subjected to enzymatic digestion in Dispase II (2.4 IU/ml) at 4 °C overnight, followed by an additional 2 h incubation at 37 °C with collagenase type I (1 mg/ml) in Eagle’s Minimum Essential Medium (EMEM) supplemented with 10% fetal bovine serum (FBS). The released cells were pelleted by centrifugation and further dissociated into single cells using 0.25% trypsin–EDTA for 20 min at 37 °C, with gentle intermittent pipetting to facilitate single-cell suspension. The reaction was quenched by washing once with EMEM containing 10% FBS, followed by two washes with sterile Ca²L/Mg²L-free PBS containing 0.04% bovine serum albumin (BSA). Cells were finally resuspended in 30–40 μl of PBS with 0.04% BSA and counted to obtain a concentration of 700–1200 cells/μl. For SC, cells were loaded onto the Chromium Controller (10x Genomics) and libraries were prepared using the Chromium Single Cell 3′ Library & Gel Bead Kit v3 (10x Genomics). Sequencing was performed on an Illumina NextSeq 2000 platform at an average depth of 30000 reads per cell (NextSeq 2000 P3 Cartridge - 100 Cycles - 1.2 billion reads distributed to 40000 cells).

### Spatial transcriptomics (ST)

Freshly enucleated whole eyes were briefly washed with PBS and fixed with 4% paraformaldehyde in PBS, 3 hours at room temperature. After fixation, eyes were washed with PBS and stored in 70% EtOH at 4 °C until tissue processing and embedding into paraffin to obtain formalin-fixed, paraffin-embedded (FFPE) tissue blocks. Blocks were sectioned in a Leica Microm HM355S rotary microtome at a thickness of 5 μm. 120 sections from before and after the target tissue depth were collected in microcentrifuge tubes and served as proxies for RNA quality control. FFPE-RNA was isolated with PureLink™ FFPE RNA Isolation Kit (ThermoFisher), following protocol instructions. The quality of isolated RNA was assessed with TapeStation (Agilent Technologies) capillary electrophoresis. Sections of the tissue target depth were collected on Epredia™ SuperFrost Plus™ Adhesion slides and air-dried, after which they were kept in a desiccated container until further analysis. High-quality samples with DV_200_ >70% were selected for analysis using Visium HD Spatial Gene Expression workflow (10x Genomics).

All downstream procedures of staining, imaging, CytAssist transfer, library preparation, and sequencing, were performed at the SciLifeLab National Genomics Infrastructure (NGI), Sweden. The sections were H&E-stained and imaged in a MetaSystems VSlide scanner, using the MetaCyte/Metafer acquisition platform with a 20x objective. Afterwards, tissues were treated with Visium HD panels and the probes were transferred onto 2×2 μm-barcoded Visium HD slides using the 10x Genomics CytAssist instrument, according to the standard Visium HD protocol. Library preparation was performed for each tissue section, and four primary libraries (corresponding to four analyzed tissue sections) were sequenced on an Illumina NovaSeq X Plus instrument (NovaSeq X Series Control Software v1.2.2.48004) using a 1.5B read flow cell in two lanes with a 43–10–10–50 read setup (read 1: 43 bp; i7: 10 bp; i5: 10 bp; read 2: 50 bp). Sequencing yielded approximately 400-600 million reads per library, with >97.9% bases ≥Q30.

Sequencing data were processed using Spaceranger v3.1.2 (10x Genomics). Raw BCL files from the NovaSeq X Plus run were demultiplexed with spaceranger mkfastq, and each of the four libraries was processed independently with spaceranger, aligned to the mm10-2020-A reference transcriptome with the STAR aligner. For each library, Spaceranger produced among other outputs, a tissue_positions.parquet file and a filtered_feature_bc_matrix.h5 file representing the full resolution 2×2μm spatial barcode grid. Standard Loupe Browser output files were also generated but were not used in downstream analyses.

### True single-cell segmentation of ST data

We developed a novel manual tracing-binning approach to achieve true single-cell resolution of ST data for the corneal epithelium (Fig. 1a-e). The approach consisted of first using ImageJ software^18^ to identify and trace individual corneal epithelial cells from the high-resolution H&E images obtained during Visium HD workflow. Next, ImageJ ROI geometries were converted into Shapely geometries in Python and further processed using custom code to resolve overlaps, self-intersections and other defects, while retaining full coverage of the tissue area. Each geometry was treated as a cell, and then a spatial matching was performed, aggregating and assigning all reads with spatial coordinates within the boundaries of each geometry, to that specific geometry (cell). Prior to aggregation, a visual inspection of the drawn geometries and the coordinates of the reads was performed, to minimize cases (<1%) where data points exactly fell on a cell boundary line. On rare ambiguous occasions, the first cell to be processed was assigned the reads and not the subsequent ones, to avoid data duplication. This produced a Scanpy anndata object with cells (observations; adata.obs) that could be treated with standard single-cell analysis tools, while for each cell we also retained information on its spatial location on a corresponding geodataframe.

**Figure 1.**
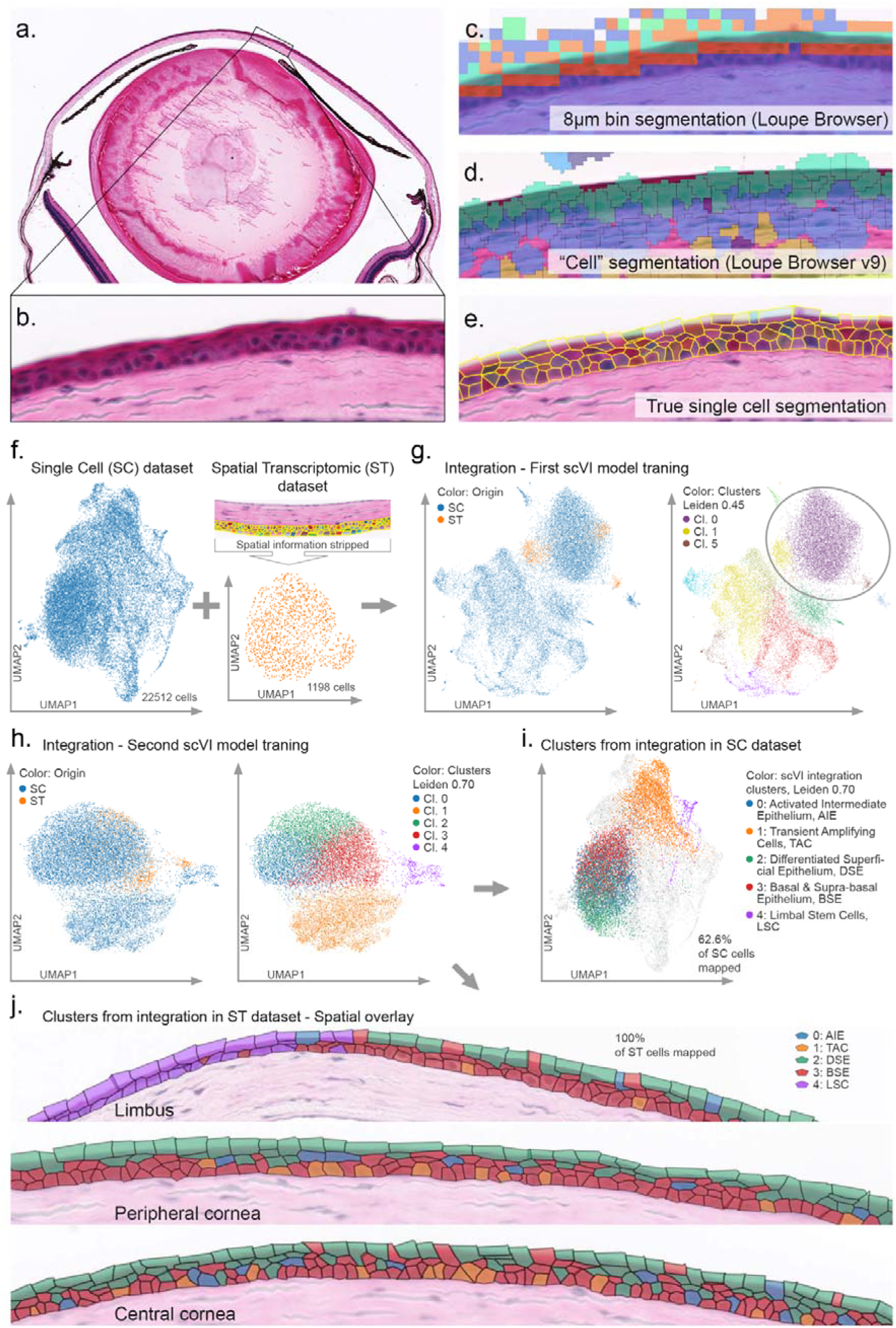
True single-cell-resolution ST integrated with SC data. **a)** H&E-stained mouse whole eye tissue section from Visium HD pipeline using MetaSystems VSlide slide scanner under a 20x objective lens. **b)** Enlarged portion of the whole-eye microscopy scan. **c)** Default “Single-cell” scale resolution with 8×8 binning from SpaceRanger visualized in Loupe browser on the same sample. **d)** Automatic cell segmentation with SpaceRanger - Loupe browser v9. **e)** True single-cell segmentation of the sample area that was used for read binning and downstream analysis. **f)** UMAP plots of full SC data from age-matched mouse corneas, n=6 and the ST cells as single-cell objects. **g)** Integration of ST and SC data through scVI model training and identification of overlapping clusters comprising cells from both datasets (clusters 0, 1, and 3). **h)** Second scVI model training performed for the isolated overlapping clusters and final clustering. **i)** UMAP plot of the SC data with post-integration cluster colors. **j)** Spatial mapping of the post-integration clusters on limbal, peripheral and central corneal epithelium in the ST subset of integrated cells.

### Quality control (QC)

Standard quality control pipelines and thresholds were utilized to detect duplets in SC data (not needed in ST data; all cells are singlets by design), to cutoff high mitochondrial content cells in SC data, and to cutoff low count cells in both SC and ST datasets. Specifically, putative doublets were assessed using Scrublet (expected doublet rate of 7.6%). Scrublet’s automated thresholding yielded a near-zero predicted doublet rate and low detectable doublet fraction, and mapping doublet scores onto the UMAP embedding did not reveal doublet enriched clusters. Based on these results, cells were retained for downstream analyses. For SC data, the mitochondrial gene content threshold was set to <15%. SC cells with less than 1000 distinctly identified genes were excluded, and subsequently cells with less than 2000 total counts were excluded. For ST cells, we used a gene count of <50 as a threshold for filtering. In SC data, genes detected in <3 cells were also filtered out, yielding an effective number of detected genes at 27,445. For additional ST samples from a 4-month-old littermate and two 2-month-old mice used for validation, the same QC criteria were applied to samples individually.

### Enrichment of ST true single-cell data through SC integration

The ST sample yielded 1201 individually traced single epithelial cells before QC filtering, while after filtering, a total of 1198 cells remained and were considered in the integration process together with 22512 QC-passed cells from the SC data at the same age (Fig. 1f). Both datasets were origin-annotated to allow for later demultiplexing and concatenated into a single anndata object, preserving all layers, and obs/vars columns by union. Then, an scVI model ^17^ was constructed, where we passed the combined anndata object, the counts layer and specified the origin as a batch key. Then the model was trained for 400 epochs, and the latent representation was saved and used for neighbors, UMAP calculations, and clustering (Fig. 1g). Because in the resulting 12 clusters (Leiden clustering, resolution 0.45), 100% of the ST cells overlapped with SC cells in 3 clusters and were completely absent from all the others, these three “overlap” clusters were isolated, creating a new anndata object that contained all spatial cells and only SC cells that were in these 3 clusters (14089 cells, or 62.6% of the SC cells). This new anndata object was fed to a second scVI model, that was similarly retrained, with the resulting latent representation used for new neighbors, UMAP calculations and clustering (Fig. 1h-j). Resulting clusters were isolated into separate single cell objects containing only cells from each respective cluster, and a secondary re-clustering of the resulting set was performed based on the same scVI latent representation. Sub-clustering was retained only for clusters where it produced meaningful cell subpopulations and then all cells were recombined back into a single dataset. Robustness of subclusters in the SC portion of the data against known dissociation-induced stress genes was evaluated by scoring the cells with the score_genes algorithm using a public gene list of 512 known dissociation-induced stress genes that commonly affect single cell studies ^19^, yielding a homogeneous score distribution across clusters with no sign of dissociation-induced stress driving clustering.

### Identification of Differently Expressed Genes (DEGs)

Differentially expressed genes were identified using Scanpy’s standard single-cell workflow. First, raw counts were normalized to a total depth of 10000 counts per cell and log transformed. Cluster designations, corresponding to the clustering scheme used for each dataset that is described in the Results section for each specific case, were used as the grouping variable for all differential expression analyses. Per cluster DEGs were computed using sc.tl.rank_genes_groups with the Wilcoxon rank sum test, without relying on the raw count layer. Visualization of the top markers was performed using Scanpy’s dot plot drawing function.

To obtain detailed per cluster gene metrics, we extracted for each cluster the ranked gene list produced by Scanpy and computed additional statistics directly from the expression matrix, including mean expression among in-cluster cells, percentage of expressing cells within (pct.1) and outside (pct.2) the cluster, and expression directionality based on log-fold change. These values were aggregated across all clusters into a single table and exported for downstream analyses. A significance threshold of adjusted p < 0.05 was applied, and cluster-specific DEGs were ordered according to their test statistic (Wilcoxon Z-scores), from highest to lowest.

### Pearson correlation for cluster similarity assessment

A Pearson correlation approach was undertaken to assess the degree of transcriptomic similarity between cell clusters within the integrated data. Pearson correlation coefficients (*r*-values) were calculated using the Z-scores for full gene lists of each cluster.

### Gene Ontology Enrichment analysis

Gene Ontology (GO) enrichment analysis for selected gene sets was performed using the Database for Annotation, Visualization and Integrated Discovery (DAVID) web service ^20,21^. Main cluster DEGs were filtered based on a threshold of p < 0.05 and absolute fold change ≥ 0.5. The top 150 genes were submitted for functional annotation. Enriched biological processes and KEGG pathways were identified, applying a false discovery rate (FDR) threshold of < 0.05 and for discovery of ST-enhanced biological processes an unadjusted P-value of < 0.05.

## Results

### Integration of scRNA-seq with custom spatial segmentation enables transcriptomic cluster enrichment and biologically accurate spatial mapping

Gene expression data from individually prepared scRNA-seq (SC) and spatial transcriptomic (ST) datasets from wild-type corneas of 4-month-old littermates were used for integration (referred to as 4mo samples hereafter). From the whole-eye histology image of the H&E-stained 4mo ST sample (Fig. 1a), the Visium HD pipeline overlay of cell geometries (Fig. 1b) illustrates the standard 8×8 µm pixelated data displayed in the default 10x Genomics Loupe Browser (Fig. 1c). Version 9 of Loupe browser from 10X Genomics supports segmentation based on automated cell detection from SpaceRanger (Fig. 1d) but does not accurately detect individual cells for true single-cell resolution. Our custom segmentation approach, however, demonstrates true single-cell spatial resolution, accurately defining cell borders based on the histology image (Fig. 1e). Integrating SC data with true single-cell-segmented ST data into a single *anndata* object while keeping track of cell origin (Fig. 1f), followed by a first scVI model training, revealed a subset of three clusters where SC and ST cells overlapped (Fig. 1g). Isolating this subset of clusters and re-training a second scVI model yielded an integrated dataset of fully overlapping SC and ST cells (Fig. 1h). Fully SC-originating clusters from the initial scVI model training that showed no overlap with ST data were excluded. Thus, in the final clustering of the integrated data, all ST-derived cells (based on histology tracing of epithelium only) were retained whereas only the SC-derived cells (from pooled whole corneas) showing clustering overlap with ST cells were retained, representing 62.6% of the full SC dataset, as visualized overlaid on the original SC UMAP (Fig. 1i).

The integrated dataset yielded biologically meaningful clustering at Leiden 0.70 resolution, without overly subdividing cells or under-representing biological variation. Importantly, all five resulting clusters contained cells from both datasets (Table S1). Up to this point, the clustering process was blinded to spatial information, but when the post-integration clusters (ST-derived cell objects) were overlaid on the histological image of the ST sample, they remarkably aligned to spatial compartments, clearly distinguishing limbal and stratified epithelial cell populations (Fig. 1j). This revealed transcriptomic homogeneity in some epithelial locations and heterogeneity in others. The integration-derived clusters (discussed in detail in the next chapter) were characterized, identified and annotated based on whole-transcriptome data with spatial visual confirmation, and are hereafter referred to as Limbal Stem Cells (LSC), Transient Amplifying Cells (TAC), Basal & Suprabasal Epithelium (BSE), Activated Intermediate Epithelium (AIE), and Differentiated Superficial Epithelium (DSE).

Overall, integrating the SC dataset with ST cells resulted in a 3.3x to 63.5x relative increase in cluster size, cell-wise, while providing a 5.9x to 17.7x increase in per-cell transcriptomic data depth, as calculated by cluster-wise average gene counts (Table S1). The number of total detected genes in the ST data portion was 19059, whereas in the SC data portion it was 27445 (44% increase), thus providing a richer pool of genes combined with deeper data, with 65.8 million reads from a sample of 6 corneas in SC, versus 0.7 million reads per corneal section in ST data.

### Integrated clusters recapitulate the full spatially resolved differentiation trajectory of corneal epithelial cells

Post-integration clusters were well-defined and cohesive, with their SC data portion broadly aligning with SC unsupervised cluster partitioning at Leiden resolution 0.3 (Fig. 2a). Interestingly, AIE, DSE and BSE overlapped mostly with SC Clusters 0 and 2, while TAC aligned with SC Cluster 1, and LSC with SC Cluster 5. Consistent with this, Pearson correlation analysis demonstrated high transcriptomic similarity across AIE, DSE and BSE (*r*-values > 0.6; Fig. 2b). Cluster gene expression profiles based on canonical markers are shown in Fig. 2c for SC/ST portions of the integrated data, along with their respective keratin profiling in Fig. 2d.

**Figure 2.**
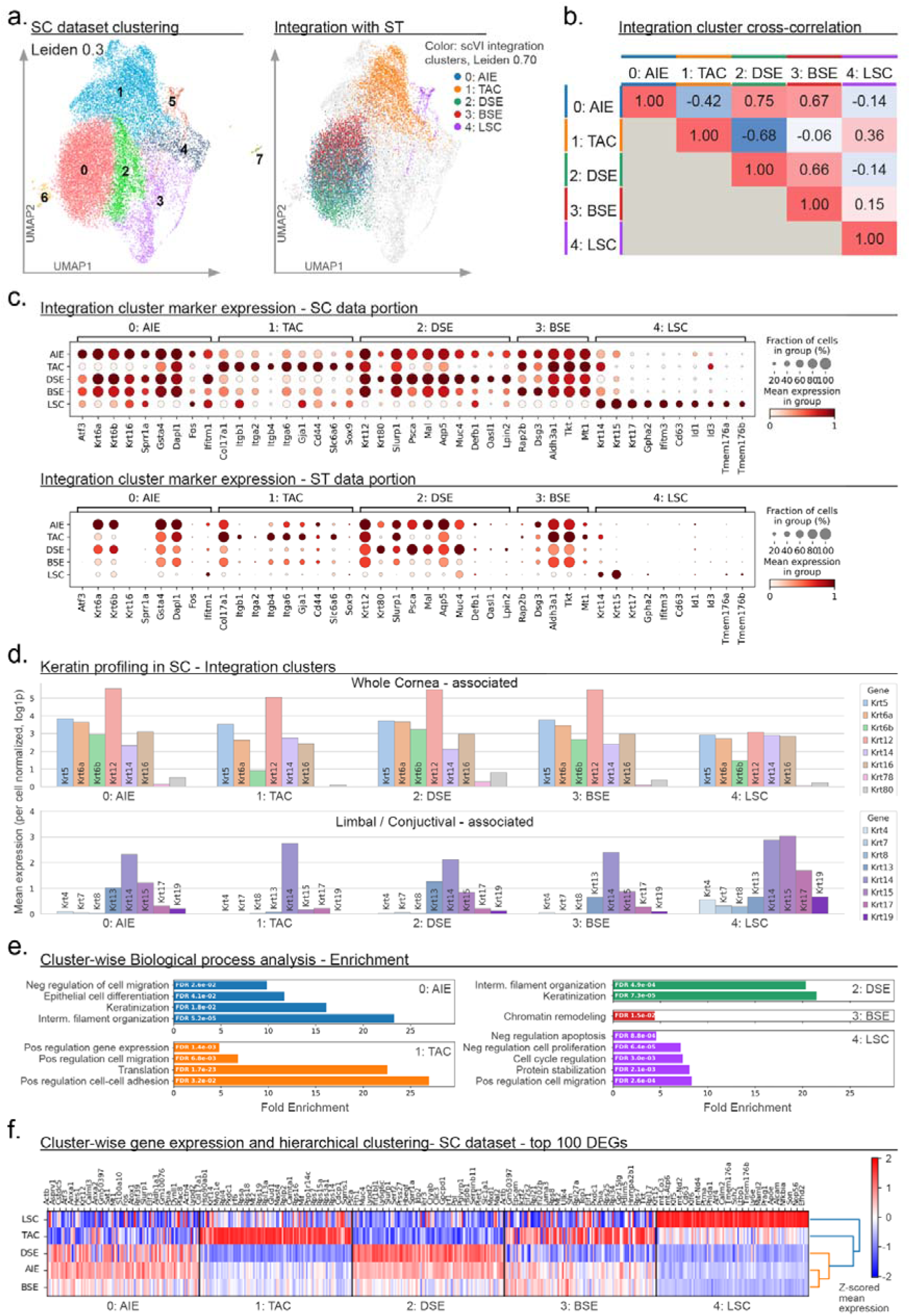
Gene expression of the integration clusters. **a)** UMAP plots of SC dataset cells with Leiden clustering at 0.3 resolution (left panel) and clustering derived from SC integration with ST cells (right panel). **b)** Pearson correlation heatmap for assessing similarity of the integrated clusters. **c)** Gene expression profile of the integration-derived clusters in the SC dataset (top panel) and the ST dataset (bottom panel). Expression values are normalized per gene between 0 (minimum average cluster expression) and 1 (maximum average cluster expression). **d)** Keratin expression profiles of the integration-derived clusters, SC data portion. **e)** Pathway analysis – enrichment of integration-derived clusters. FDR- and fold-enrichment based top significant biological processes per cluster are presented. **f)** Gene expression and hierarchical clustering based on the top 100 DEGs per cluster.

#### Limbal stem cells (LSCs)

The LSC cluster was distinctly confined to the dual-layer epithelium in the limbal zone (Fig. 1j) and was enriched in *Gpha2*, *Ifitm3, Cd63, Id1, Id3,* and *Tmem176a/b*, all LSC-associated genes^3,5,6,22^ as well as keratin patterns associated with conjunctiva (*Krt4, Krt7, Krt8, Krt19*) ^3,5,23,24^ and/or stem/progenitor cells (*Krt14, Krt15, Krt17)* ^25–28^(Fig. 2c, d). LSC also exhibited suppression of *Krt12, Krt78, Krt80, Slurp1* and *Aqp5 (*Fig. 2c), representing negative LSC markers related to further differentiation^29–32^. Spatial localization of *Krt14, Krt15, Krt19* and *Gpha2* in the limbus aligned with LSC distribution (Fig. 3), with individual *Gpha2+* and *Krt15+* cells scattered in the basal and suprabasal layers extending into the central cornea (Fig. S1a), while *Krt12* and *Slurp1* declined in expression towards the limbus (Fig. S1b). GO analysis revealed that LSC was enriched in processes consistent with homeostasis (Fig. 2e, Table S2), including negative regulation of proliferation and apoptosis, cell cycle regulation, and enrichment of cell migration and endocytosis suggesting priming for epithelial repair, while responsiveness to hypoxia and oxidative stress highlights an adaptive stress response of LSC promoting long-term survival and maintenance over acute stress-related differentiation.

**Figure 3.**
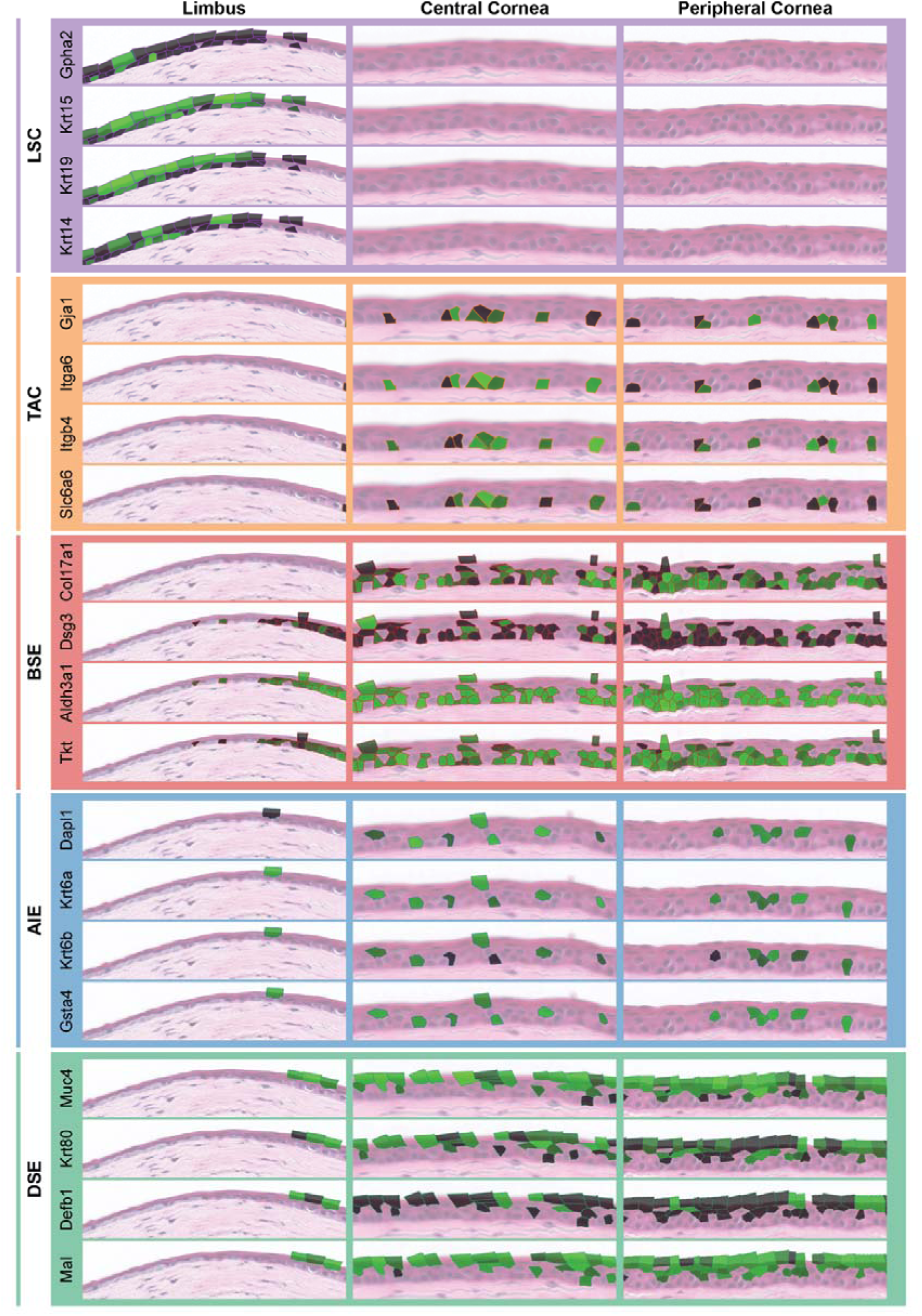
Spatial gene expression maps of cluster-specific markers across the epithelium. Only cells comprising the cluster in question are mapped for clarity of cluster visualization. Intensity of green shading of cells signifies per-gene normalized expression level, with maximum/minimum intensity representing log-transformed maximum/minimum expression across all cells in the cluster.

#### Transient Amplifying Cells (TAC)

The TAC cluster was confined to the basal layer in peripheral and central regions, in a spatially sparse, non-contiguous manner (Fig. 1j). Accordingly, TAC exhibited strong canonical basal cell signature enriched for various integrins (*Itga2, Itga6, Itgb1, Itgb4*)*, Col17a1, Cd44* and *Gja1,* which serve as mediators of cellular adhesion, extracellular matrix crosstalk and cell-to-cell communication in basal cells ^33–36^(Fig. 2c). TAC was additionally enriched in *Slc6a6* and *Sox9* (Fig. 2c), both shown to have important functional roles in corneal progenitors ^4,37^, and canonical proliferation markers (*Mki67, Top2a, Birc5, Pcna*) (Fig. 4a). The TAC keratin profile was enriched in progenitor-associated *Krt14*, while differentiation- and limbal-associated keratins were downregulated (Fig. 2d). Spatial mapping of *Itgb1, Itga6, Slc6a6* and *Gja1* demonstrated the expression of these markers in the TAC cluster across the peripheral and central corneal epithelium (Fig. 3). TACs were enriched for translational and ribosomal biogenesis programs indicative of high protein synthesis capacity, cell adhesion and hemidesmosome assembly pathways, as well as positive regulation of cell migration, all hallmarks of a highly active, rapidly proliferating and centripetally migrating transiently amplifying population (Fig. 2e, Table S2).

**Figure 4.**
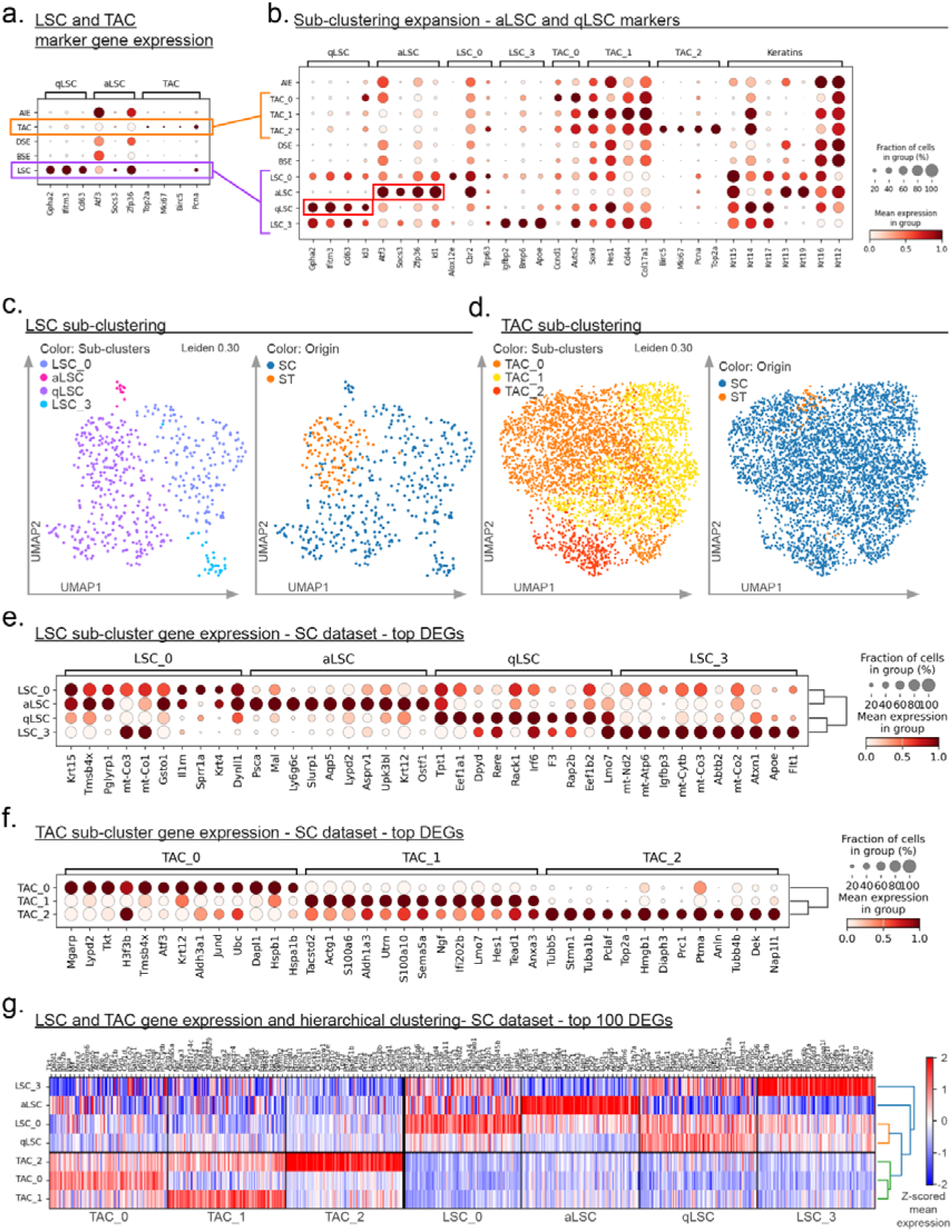
LSC and TAC sub-clustering for finer fidelity cell type sub-population identification. **a)** Distribution of putative qLSC, aLSC and canonical TAC markers across clusters. **b)** Gene expression profile for sub-cluster-defining genes and keratins. **c)** LSC UMAP projection showing sub-cluster segmentation at resolution Leiden 0.30 (left panel) and the distribution of SC and ST single cells across the same UMAP coordinates (right panel). **d)** TAC UMAP projection with sub-cluster segmentation at resolution Leiden 0.30 (left panel) and the distribution of SC and ST single cells across the same UMAP coordinates (right panel). **e)** Dot plot of the top 10 DEGs of the LSC sub-clusters. **f)** Dot plot of the top 10 DEGs of the TAC sub-clusters. **g)** Sub-cluster hierarchy heat map for 100 genes per sub-cluster. The dot plot expression values are normalized and scaled per gene between 0 (minimum average cluster expression) and 1 (maximum average cluster expression).

#### Basal and Suprabasal Epithelium (BSE)

The BSE cluster represented the largest proportion of ST cells and localized to basal and suprabasal layers across the entire epithelium (Fig. 1j, Table S1). Consistent with this spatial pattern, BSE was enriched in genes overlapping with basally located TAC, including corneal crystallins *Aldh3a1* and *Tkt* as well as *Col17a1* (Fig. 2c), and the keratin profile showed suppression of *Krt13, Krt78,* and *Krt80* (Fig. 2d). In contrast to TAC, however, BSE did not express proliferation markers (Fig. 4a, b). Spatial mapping of shared TAC/BSE markers *Aldh3a1, Tkt* and *Col17a1* demonstrated strong enrichment of these markers in basal and suprabasal cells assigned to BSE (Fig. 3). The only significantly enriched process in the BSE was chromatin remodeling (Fig. 2e, Table S2), consistent with epigenetic reprogramming underlying epithelial stratification and differentiation.

#### Activated Intermediate Epithelium (AIE)

The AIE cluster comprised a population of cells distributed across the suprabasal limbus and basal and suprabasal peripheral and central corneal epithelium (Fig. 1j) and exhibited an intermediate spatial and transcriptomic profile between BSE and DSE clusters (Fig. 2a, b, f). AIE was enriched for *Dapl1,* an early epithelial differentiation activation marker ^38^, *Atf3, Gsta4* and *Fos,* which are known for their roles in stress-responses^14,39^, as well as *Sprr1a* and *Ifitm1,* which have been shown to participate in regenerative programs^40,41^ (Fig. 2c). The AIE keratin profile was closely aligned with that of BSE and DSE, expressing *Krt6a/b* and *Krt16*, keratins known to be induced in epithelial activation during stress and regeneration ^42,43^, although they are also present in homeostatic situation^27^ (Fig. 2c, d). Higher *Krt12* expression coupled with lower *Krt14* indicates differentiation. Spatial mapping of *Gsta4, Dapl1*, and *Krt6a/b* (Fig. 3) confirmed their expression in epithelial layers where AIE cells reside. AIE was enriched in intermediate filament organization and keratinization processes, indicating cytoskeletal and keratin network remodeling associated with epithelial maturation. Co-enrichment of epithelial differentiation along with suppression of migratory programs indicates differentiation-associated stabilization (Fig. 2e, Table S2).

#### Differentiated Superficial Epithelium (DSE)

The DSE cluster populated the most anterior layers of the central and peripheral corneal epithelium (Fig. 1j) and was enriched for *Psca, Mal, Slurp1, Aqp5* and *Muc4,* genes associated with the superficial corneal epithelium ^10^. Moreover, DSE expressed *Defb1*, a beta defensin indicated in innate and inflammatory signaling responses ^44^; *Oasl1,* an classical interferon-stimulated gene^45^ and *Lpin2,* another regulator of interferon-mediated immune responses ^46^ (Fig. 2c). DSE exhibited upregulation of *Krt12, Krt13, Krt78,* and *Krt80*, along with *Krt6b* (Fig. 2d). Spatial mapping of *Krt80, Muc4, Mal* and *Defb1* (Fig. 3) confirmed strong expression of these markers in the most superficial cells. DSE was enriched for processes of keratinization and intermediate filament organization, consistent with terminal epithelial differentiation characterized by stabilized keratin networks and structural maintenance (Fig. 2e, Table S2).

### Transcriptional relationships underlying corneal epithelial differentiation

Together, the five integrated clusters represent the complete differentiation trajectory of the corneal epithelium, from LSC to TAC, BSE, AIE and finally to DSE, with annotations strongly supported by integrated data and spatially resolved gene expression patterns at the individual cell level. A heatmap depicting the top 100 marker genes for each cluster (Fig. 2f) highlights distinct transcriptional profiles and lineage-related similarities, particularly between LSC-TAC, TAC-BSE and BSE-AIE-DSE. Notably, AIE had similar transcriptomic expression to BSE and DSE, consistent with its role as a transitional cell state completing the epithelial differentiation trajectory.

### Detailed investigation of cell sub-populations

We next used the increased transcriptomic resolution afforded by SC-enrichment to resolve potential epithelial subpopulations within main clusters of interest (Fig. 4a). First considering stem and progenitor clusters, sub-clustering based on full transcriptomic profiles yielded four LSC (LSC_0-3) and three TAC (TAC_0-2) sub-clusters (Fig. 4b). UMAP depiction of SC and ST single cells across distinct sub-clusters indicate ST/SC contributions (Fig. 4c,d).

All LSC sub-clusters expressed limbal *Krt15* (Fig. 4b). We identified reported signatures of quiescent (qLSC; *Gpha2, Ifitm3, Cd63*) and activated (aLSC; *Atf3, Socs3, Zfp36*) sub-clusters ^3^ to annotate LSC_1 and LSC_2 as aLSC and qLSC, respectively (Fig. 4b, e). qLSC was enriched with *Id3,* an inhibitor of differentiation ^47^, along with basal/progenitor keratins *Krt14* and *Krt17* while aLSC was enriched in *Id1, Krt4, Krt13,* and *Krt19* (Fig. 4b, e). Top aLSC DEGs overlapped with differentiation-associated markers enriched in DSE (Fig. 2d), including *Psca, Mal, Slurp1* and *Krt12* (Fig. 4e). Sub-cluster LSC_0 had an intermediate qLSC/aLSC marker profile, accompanied by expression of LSC hallmark gene *Trp63* along with *Krt4, Alox12e,* and *Cbr2*, which are predominantly expressed in conjunctival epithelium ^43^. LSC_3 was qLSC-like, expressing *Gpha2, Ifitm3* and *Cd63* along with additional putative qLSC markers *Apoe* and *Igfbp2* ^6^ (Fig. 4b, e). Notably, only the qLSC sub-cluster overlapped with the ST portion of the data, apart from a single ST cell object in LSC_0 (Fig. 4c).

TAC_0 was characterized by *Mgarp, Lypd2, Tkt, Aldh3a1, Dapl1* and *Ccnd1* a cell cycle regulating cyclin ^48^, and exhibited the strongest *Atf3* expression within the TAC subclusters, concomitant with increased *Krt12/Krt14* expression ratio (Fig. 4b, f). By contrast, TAC_1 was enriched in *Col17a1, Cd44, Sox9* and *Hes1*. Finally, TAC_2 was strongly enriched with proliferation markers *Top2a, Mki67, Birc5, Pcna* and *Trp63*, with top 10 DEGs further including additional TAC-associated markers *Stmn1*, *Pclaf, Tubb5, Tuba1c and Prc1*^6^ (Fig. 4b, f). Of TAC sub-clusters, TAC_1 was exclusively represented in the SC portion, while the ST portion overlapped predominantly with TAC_0 (Fig. 4d).

The heatmap and sub-clustering hierarchy (Fig. 4g) distinguishes LSC from TAC populations, reflects the common origin of TAC subclusters, and highlights distinct gene sets defining aLSC and LSC_3 populations.

Given the large BSE population represented across multiple epithelial layers, we additionally sub-clustered BSE in the same manner as above, yielding three subpopulations (BSE_0-2) that were distributed across both SC and ST data in the UMAP (Fig. 5a). Sub-clusters exhibited distinct profiles based on selected markers of stemness, differentiation, and adhesion, as well as top DEGs (Fig. 5b). BSE_0 is distinguished by *Gsta4*, *Lypd2*, and *Tmsb4x* expression, exhibiting the highest expression of *Atf3* and *Zfp36*, along with intermediate *Krt12* and high *Krt15* expression. BSE_1 expresses *Krt6b*, *Dsg3* and *Muc4* along with the highest differentiation-associated *Krt12* and lowest *Krt15*. Finally, BSE_2, comprising 64% of cells in BSE, is enriched in *Dapl1*, *Tkt*, *Col17a1* and *Itga6*, also exhibiting the lowest level of *Krt12*. Based on these expression patterns alone, we hypothesized BSE_2 would localize to the anatomic basal epithelial layer in contact with the epithelial basement membrane, while BSE_0 would localize to an intermediate layer and BSE_1 to the most superficial layer of BSE. Isolating the ST portion of BSE cells and mapping spatial coordinates to the original tissue locations indeed revealed a cell localization in full accordance with this pattern (Fig. 5c).

**Figure 5.**
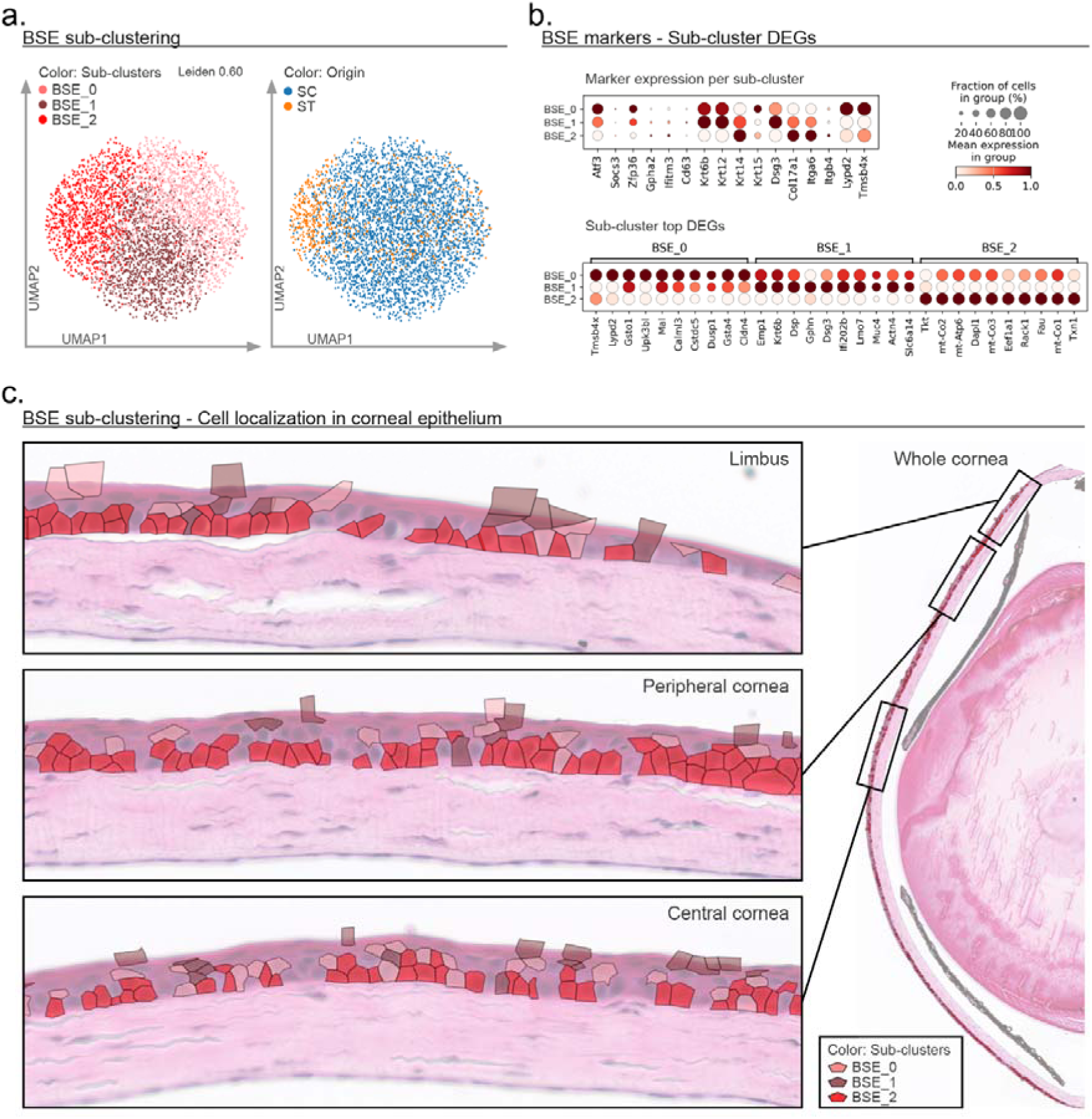
BSE sub-clustering and sub-cluster localization. **a)** BSE cells UMAP projection showing the segmentation of cells into sub-clusters (left panel) at resolution Leiden 0.60 and the distribution of SC and ST single cells in the same UMAP coordinates (right panel). **b)** Dot plot visualization of curated BSE marker expression across the three BSE sub-clusters (top panel) and the expression top 10 DEGs between sub-clusters. The dot plot expression values are normalized and scaled per gene between 0 (minimum average cluster expression) and 1 (maximum average cluster expression). **c)** Localization of the BSE cells in the corneal tissue, and distribution of sub-clusters in the limbus (top panel), peripheral cornea (middle panel) and central cornea (bottom panel).

### Genes and biological processes specific to ST data

Despite the greater depth of the SC portion, some genes were more prominent in the ST portion of the integrated dataset. A total of 43 genes were identified with a 3-fold ST/SC expression ratio and a minimum 0.5 log-transformed total expression level in the ST portion (hereafter termed ‘ST-enhanced’, Table S3, Fig. S3). Among ST-enhanced genes, oscillatory, circadian rhythm-associated ^49^ *Dbp* (ST/SC ratio: 23.6)*, Hr* encoding homeostatic epithelial Hairless protein ^50^ (ST/SC ratio: 14.9), and *Sema3f* (ST/SC ratio: 12.0) were prominent. *Fam129b* was uniquely detected in the ST data and absent in SC data. Spatial localization of ST-enhanced genes (Fig. S2) indicated limbal and abundant central corneal expression patterns. GO term enrichment analysis identified the following ST-enhanced biological processes: nervous system development (GO:0007399), axon guidance (GO:0007411), neural crest migration (GO:0001755), regulation of neuron projection development (GO:0010975), and response to wounding (GO:0009611) (Table S4). We examined whether genes contributing to spatially emphasized processes corresponding to the GO terms above indicated any cluster-specific distribution; however, no cluster specificity was observed, indicating processes represented throughout the epithelium.

### Validation and extension of ST-SC integration to new ST samples

As a validation of our true single-cell ST method and SC-ST integration approach, we applied gene expression profiles of the above described clusters from the 4-month-old WT cornea to a new, previously unanalyzed WT cornea from a different 4-month-old mouse and two additional corneas from two WT mice aged 2 months. To achieve this, ocular tissue sections from these additional corneas were first sequenced on the Visium HD platform, followed by application of our tracing pipeline to create single-cell ST objects. Overlays were then generated to visualize cell objects expressing gene signatures characteristic to our five previously identified integrated clusters (Fig. S4). Expression patterns in the independent 4-month WT cornea confirmed spatial localization of markers characteristic of each cluster. Moreover, in separate corneas at 2 months of age, expression remained consistent with cluster annotations; however, at 2 months elevated gene expression in the limbus was evident.

### AIE cluster identification in human public datasets

To validate the AIE cluster, we investigated the publicly available Human Corneal Cell State Meta Atlas dataset ^51^ against AIE signature genes (Fig. S5), by calculating a per-cell score, based on the top 20 AIE DEGs (scanpy score_genes function). Interestingly, this scoring revealed an unusually high and specific correlation with a cluster annotated as “5: Cj”. This cluster, as our AIE cluster, represents a major cell population (10198 cells in the meta-dataset), with a distinct signature of low *KRT14* and high *KRT12*, along with expression of *KRT13*, *KRT15*, and *AQP5*, all of which signify a differentiating state and mirror our AIE cluster expression pattern (Fig. 2c, d). In the meta-atlas, the “5: Cj” cluster was adjacent to, and partially overlapped with, the “8: Wing CE” cluster in the UMAP (Fig. S5), indicating its similarity to wing cells but notably differentiated by absent *KRT13*, *KRT15* and *AQP5* expression in the wing cell cluster as are currently known as conjunctival gene signature. Without the benefit of ST, the human atlas annotation cannot be localized within the tissue; however, strong overlap with the present dataset suggests a high similarity of “5: Cj” with our AIE cluster and its non-conjunctival localization.

## Discussion

Here, we present an integrated, spatially accurate true single-cell dataset defining the healthy adult mouse corneal epithelium, combining Visium HD ST with matched scRNA-seq data, both based on 10x Genomics platforms. To address an existing gap in integration approaches for high-resolution transcriptome-wide SC/ST data ^52^, we developed a custom true single-cell segmentation for ST enabling spatial and single-cell objects to be treated consistently across both platforms. Remarkably, following data integration, we were able not only to assign and map all clusters to biologically relevant epithelial compartments within the murine cornea, representing the full differentiation trajectory from LSC to mature superficial epithelium, but also to gain deeper biological insights at the subcluster level and to identify a activated signature mostly highlighted as a new compartment located between BSE and DSE clusters named AIE, revealing additional insights into the epithelial differentiation trajectory.

LSC localized to the bilayer limbal region across postnatal ages 2 – 4 months. qLSC and putative aLSC signatures were detected, with qLSC consistent with reported markers *Gpha2, Ifitm3, Cd63, Krt14, Krt17* and *Id3* ^3,5,6^. An aLSC population previously proposed ^3^ but not uniformly identified in a subsequent study ^5^ was nonetheless detected in our data as *Atf3/Socs3/Zfp36* putative aLSC with additional upregulation of limbal keratins *Krt15* and *Krt19,* along with *Id1* known to inhibit lineage commitment and regulate quiescence in stem cells ^53,54^, in agreement with a recently reported *Atf3+/Id1+* limbal sub-cluster ^7^. Concomitant upregulation of differentiation-associated genes in our data including *Slurp1, Krt12,* and *Krt13* suggests differentiation-primed aLSC. Notably, *Apoe* in our LSC_3 subcluster is a recently proposed LSC marker ^22,43,55^ reported in a qLSC-annotated subcluster ^6^. These results are in part functionally supported by *Apoe*-null mice developing corneal epithelial defects ^56^. Here, we identify *Apoe-*expressing LSC_3 distinct from qLSC and aLSC subpopulations, suggesting LSC_3 may represent a unique LSC transcriptional profile warranting further investigation.

Our TAC localized to the basal layer with subclustering resolving three distinct subpopulations. TAC_2 has a proliferative gene signature (*Top2a, Mki67, Birc5, Pcna* and *Trp63)* consistent with classical TAC, whereas TAC_1 enriched for *Sox9* and *Hes1* suggests an early TAC state, supported by *Sox9* maintaining clonogenic and normal differentiation potential and *Hes1* inhibiting the differentiation of corneal progenitors ^57–59^. Lastly, TAC_0 is enriched with *Ccnd1* and exhibits an activated-differentiating signature with increased *Atf3*, *Dapl1,* and *Krt12* expression. *Ccnd1* plays a central role in progenitor cell expansion and inhibition of differentiation ^60^ and indicates retained proliferative capacity, with *Dapl1* being expressed in early differentiating corneal epithelial cells ^38^, together suggesting a mature, differentiation-committed TAC_0. Our findings align with scRNA-seq identification of three murine TAC subpopulations differing in maturation and proliferative status ^5^.

Differentiation-primed, activated subclusters were identified across LSC, TAC, and BSE, characterized by elevated *Atf3* expression and most prominently observed in AIE. Strong enrichment of *Krt16, Dapl1*, *Sprr1a* and *Gsta4,* markers activated during wound healing ^38–40,43^ distinguished AIE from main BSE and DSE clusters. An activated intermediate population in the central and peripheral cornea has not been explicitly described previously, although *Atf3* expression was apparent in a putative wing cell cluster ^3^, and is known to promote growth and repair, responding to cellular stressors along with *Krt16* and growth-associated *Sprr1a* ^2,61–63^. The AIE transcriptomic signature thus suggests active differentiation/transition while protecting against cellular stress. A similar relatively enriched active differentiating signature (*Atf3, Zfp36, Gsta4,* and *Dapl1*) was detected in BSE_0 which localized to suprabasal epithelial layers – the same layers where AIE is present – supporting transcriptional similarity of BSE_0 with AIE and corroborating the localization of AIE within suprabasal/wing layers. In contrast, basally localized BSE_2 exhibits an adhesion-related gene signature, including *Col17a1*, *Itga6*, and *Itgb4*, with transcriptional similarity to TAC, whereas BSE_1 localized to more superficial epithelial layers with a clear differentiation signature and similar to DSE expresses *Muc4*.

Taken together, our data show a common activated signature across aLSC, TAC_0, AIE and BSE_0 that we hypothesize represents a shared priming of corneal epithelial cells for state transitions at multiple differentiation levels. Our findings suggest that at the transcriptomic specification level, less differentiated states pass through this activated state as they transition to more differentiated populations. The intermediate epithelium is comprised of activated epithelial states such as BSE_0 and AIE that are spatially distributed within the anatomic wing layer, together with non-activated BSE and DSE cells. The anatomic wing cell layer does not appear to represent a singular, transitional stage, nor do its boundaries define a homogeneous transitional cell state. Supporting this, we additionally found a strong similarity score of our AIE with a well-defined cluster in multiple human corneal scRNA-seq datasets^51^, which additionally had transcriptional similarity to wing cells. In the absence of an available AIE cell state description, the cluster had been loosely defined, and our data suggest a refined interpretation also relevant to human samples. Notably, *Krt13* at the transcriptional level alone is insufficient to annotate epithelial cells as ‘conjunctival’, as *Krt13* transcripts are present in non-conjunctival AIE and DSE clusters across the normal corneal epithelium. Our additional examination of AIE markers in younger 2-month-old WT mouse corneas localized these cells to the intermediate corneal epithelium to further corroborate this transcriptional state. Most published SC datasets sequencing mainly corneal tissue detect *KRT13* transcripts across a greater proportion of cells than would be expected based on the amount of conjunctival tissue present in analyzed samples. Our findings prompt a re-evaluation of published SC annotations based on *KRT13* transcripts.

It is known that dissociation of tissues for scRNA-seq disrupts native cell-cell and cell-matrix interactions^14,64^, and among ST-enhanced genes was *Fam129b* (*Niban2*) which exerts antioxidant and cytoprotective effects ^65,66^, positively regulating cell motility ^67^. *Sema3f,* enhanced 12-fold in ST, is an axon guidance cue in adult corneal epithelium downregulated in response to injury ^68,69^, that together with *Pax6* and *Sema4d* is associated with axon guidance and neural crest migration processes, suggesting corneal innervation programs of vital importance for epithelial homeostasis are preserved in ST data. Further, ocular development master regulator *Pax6* ^70^ was four-fold enhanced in ST data, suggesting the potential value of ST in decoding *Pax6*-dependent transcriptional programs, including those relevant for ocular development. Further ST-enhanced factors *Itgb4, Tns4,* and *Tjp3* highlight epithelial cell attachment, adhesion and extracellular signaling as important mechanisms preserved by avoiding cell dissociation. Integrin subunit β4, encoded by *Itgb4*, heterodimerizes with integrin subunit α6 to mediate basal epithelial cell attachment to basement membrane via hemidesmosomes ^33^, with both subunits along with *Tns4* highly expressed in basally located TAC and BSE clusters (Fig. 2c, Fig. 3, Fig. S2). Importantly, *Tns4* regulates extracellular matrix biomechanics and focal adhesion signaling in the mouse corneal epithelium ^71^, while *Tjp3* (also known as *zona occludens 3* or *ZO-3*) codes for an epithelial tight junction protein ^72^ which interestingly, has not previously been reported in the cornea. These findings suggest that ST may better preserve microenvironment-dependent and corneal epithelial barrier genes.

The five major epithelial cell populations we detected were remarkably stable when applied to additional WT tissues of the same age (4 months) and an earlier postnatal age (2 months). The true-single-cell integration approach may thus enable deeper analysis of ST samples lacking corresponding SC data, providing a less resource intensive SC-independent approach for further ST-based studies. We noted clusters at 4 months of age mapped to the same spatial compartments of 2-month-old mice, although gene expression was more pronounced, and extended further into, the limbal area at the earlier postnatal age, in accordance with known maturation of mouse corneal epithelium between 3 and 6 months postnatally ^73^ and highlighting the need to investigate sufficiently mature mice to decode programs of mature epithelial homeostasis.

The present study had several limitations. The transcriptomic depth of ST was lower than with SC data. Greater sequencing depth or optimization of histology section thickness could be further explored to improve ST robustness for deeper investigation and localization of subpopulations. Nevertheless, the data depth achieved by single-cell approaches, where the entire cell is profiled, is unlikely to be fully recapitulated by technologies that capture only a section of each cell. This inherent limitation presents challenges for data integration, with varying degrees of success. In addition, although we focused on gene regulatory programs of homeostasis, protein-level studies are needed for further, functional-level insights. Newer spatial proteomics technologies enable targeted multi-protein detection, and our data can inform panel selection for high-throughput analyses.

To conclude, we identified a putative AIE cell population and related activated differentiating state across TAC, LSC and BSE subpopulations that warrant further investigation. Spatial localization at the single-cell level enables specification of a heterogeneous anatomic wing cell layer consisting of differentiated and transitioning cell populations. We provide our dataset of transcriptomic data at single-cell resolution as a resource to investigate genetic programs underlying murine corneal epithelial homeostasis, highlighting that the true-single-cell ST may preserve gene expression programs not apparent in SC data, including axon-dependent signaling, cell adhesion regulation programs, and epithelial barrier creation by cell-to-cell and cell-to-extracellular matrix junction regulation.

## Acknowledgements

The authors thank Åsa Schippert and Mouna Tababi for providing guidance and assistance with NextSeq 2000 sequencing. We also wish to acknowledge the Biology Core facility at the Faculty of Medicine and Health Sciences, Linköping University, specifically the molecular biology labs and the histology lab, for the availability of infrastructure and expertise regarding the transcriptomic experiments and FFPE sample preparation – sectioning. We also thank Xesús Abalo for coordinating and facilitating the Visium HD library preparation and sequencing of our ST samples at the National Genomics Infrastructure (NGI) Sweden labs. This work was funded by the European Joint Programme on Rare Diseases (EJP-RD) under the project AAK-INSIGHT, Grant No. EJPRD20-135, and the Finnish Silmäsäätiöiden tohtoritutkijapooli (SSTTP).

## Author contributions

N.L and P.M. conceptualized and designed the study; D.J., M.V., N.L. and P.M. developed the data analysis methodology; D.J. performed surgical procedures; D.J. and P.M. performed single cell experiments, sequencing, all wet lab procedures and ST sample preparation; P.M. performed all bioinformatic pipelines, data integration, data output, statistics and visualization; D.J, M.V. and N.L. performed data interpretation, cluster annotation and marker identification; D.J. performed biological process enrichment; D.J, M.V. and N.L. wrote and edited the manuscript; N.L. and P.M. revised the manuscript; N.L. secured funding.

## Supplementary information and Data Availability

Supplementary Information consists of Supplementary Tables S1-S4 and Supplementary Figures S1-S5. The raw data supporting this study have been deposited in Zenodo (DOI: 10.5281/zenodo.20071121) and are available at https://doi.org/10.5281/zenodo.20071121.

## Supplementary materials

**Table S1.**
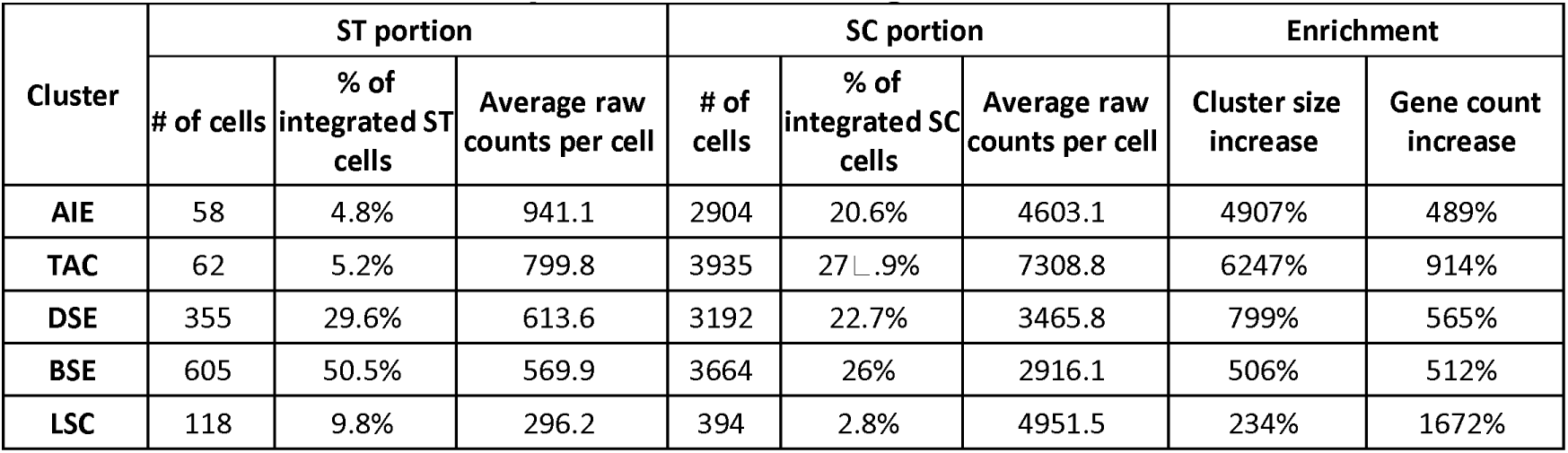
Cluster cell composition after integration.

**Table S2.**
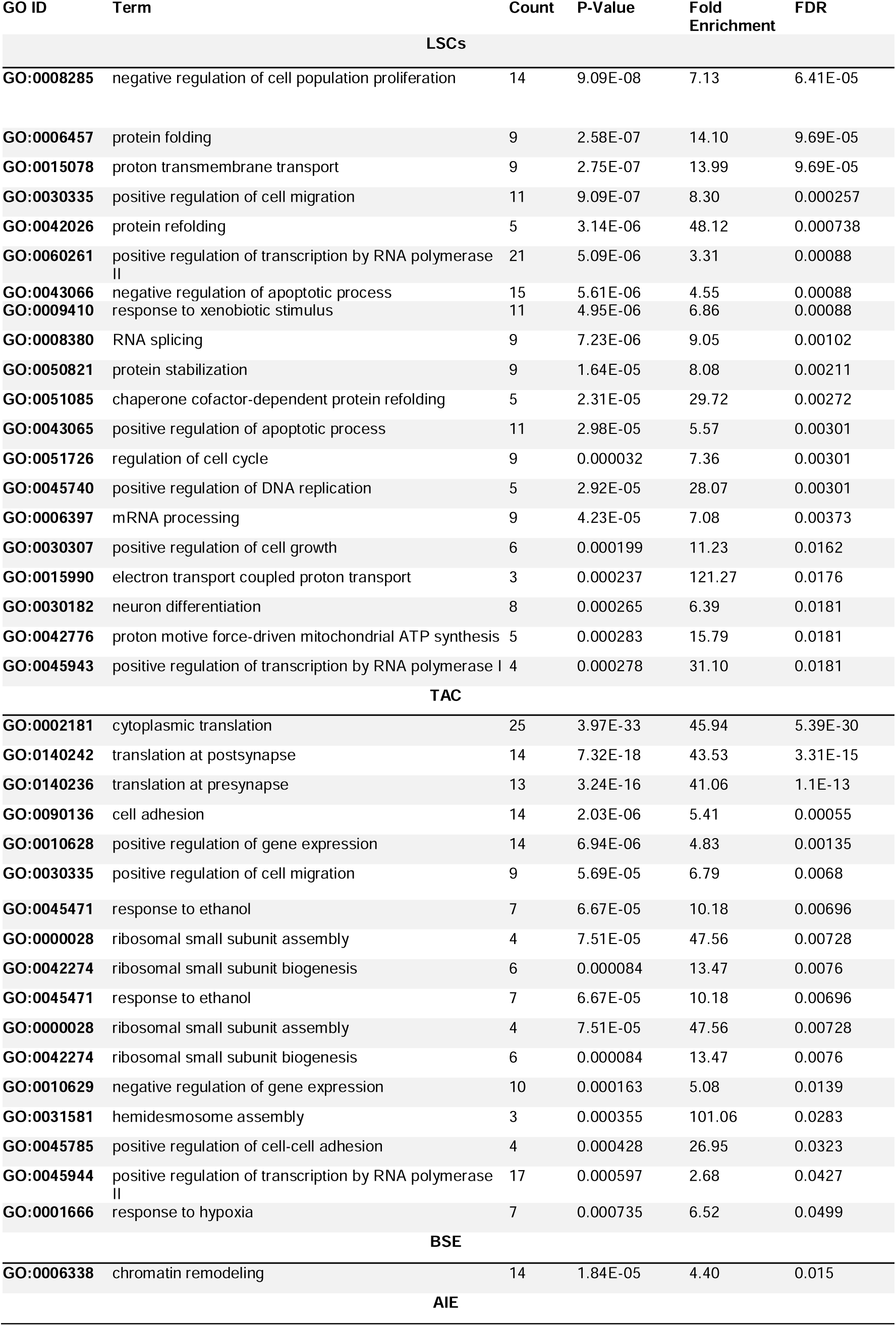
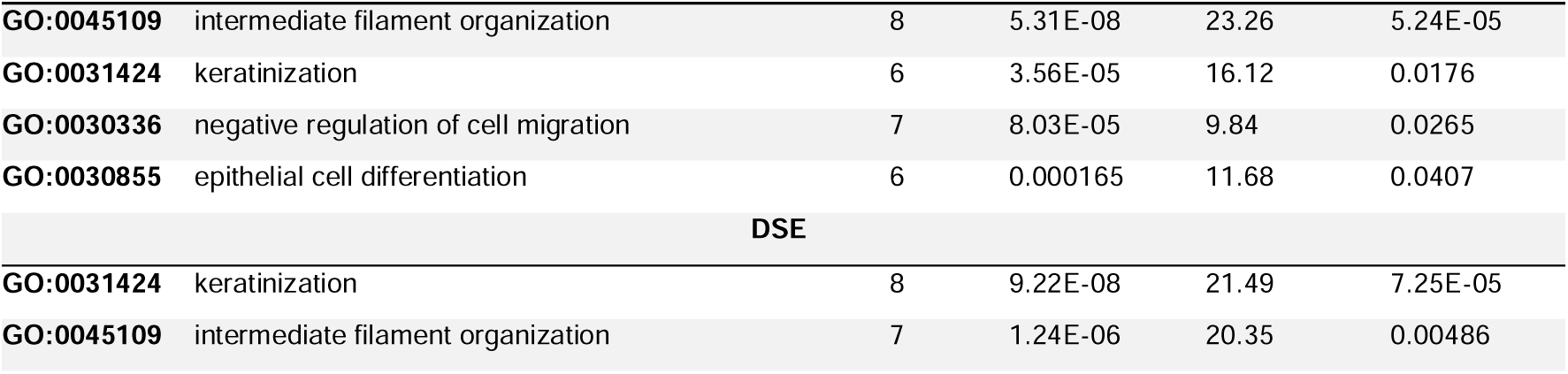
Full list of biological processes enriched across integrated clusters.

**Table S3.**
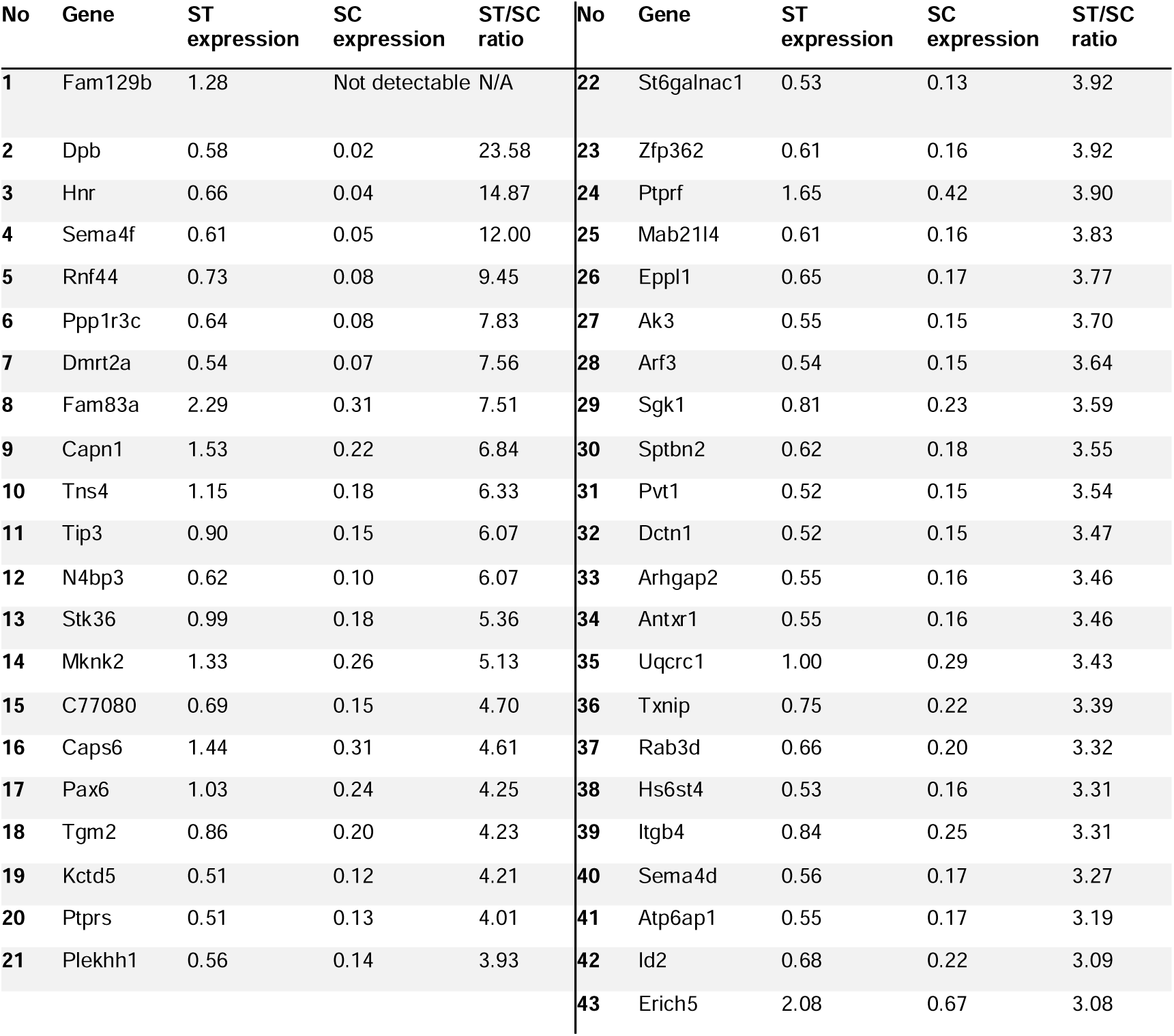
ST-enhanced genes. The full list of 43 ST-enhanced genes, based on minimum total log-normalized expression level of 0.5 and > 3-fold ST/SC expression ratio

**Table S4.**
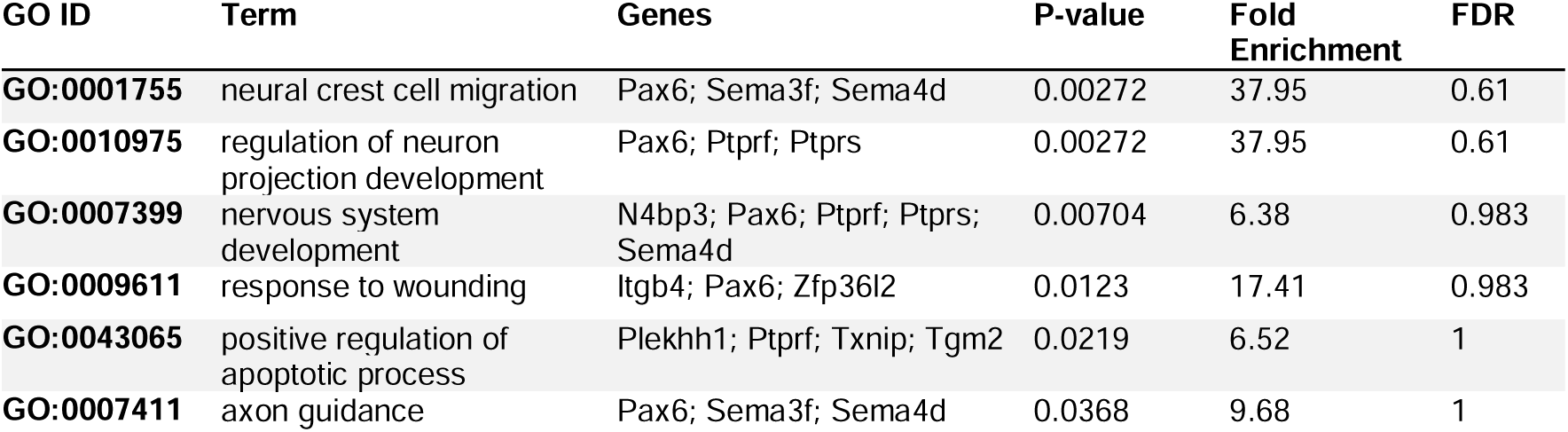
ST-enhanced biological processes. GO term enrichment of spatially emphasized biological processes, based on unadjusted p-value < 0.05 and ST-enhanced gene list and minimum count of three genes.

**Figure S1.**
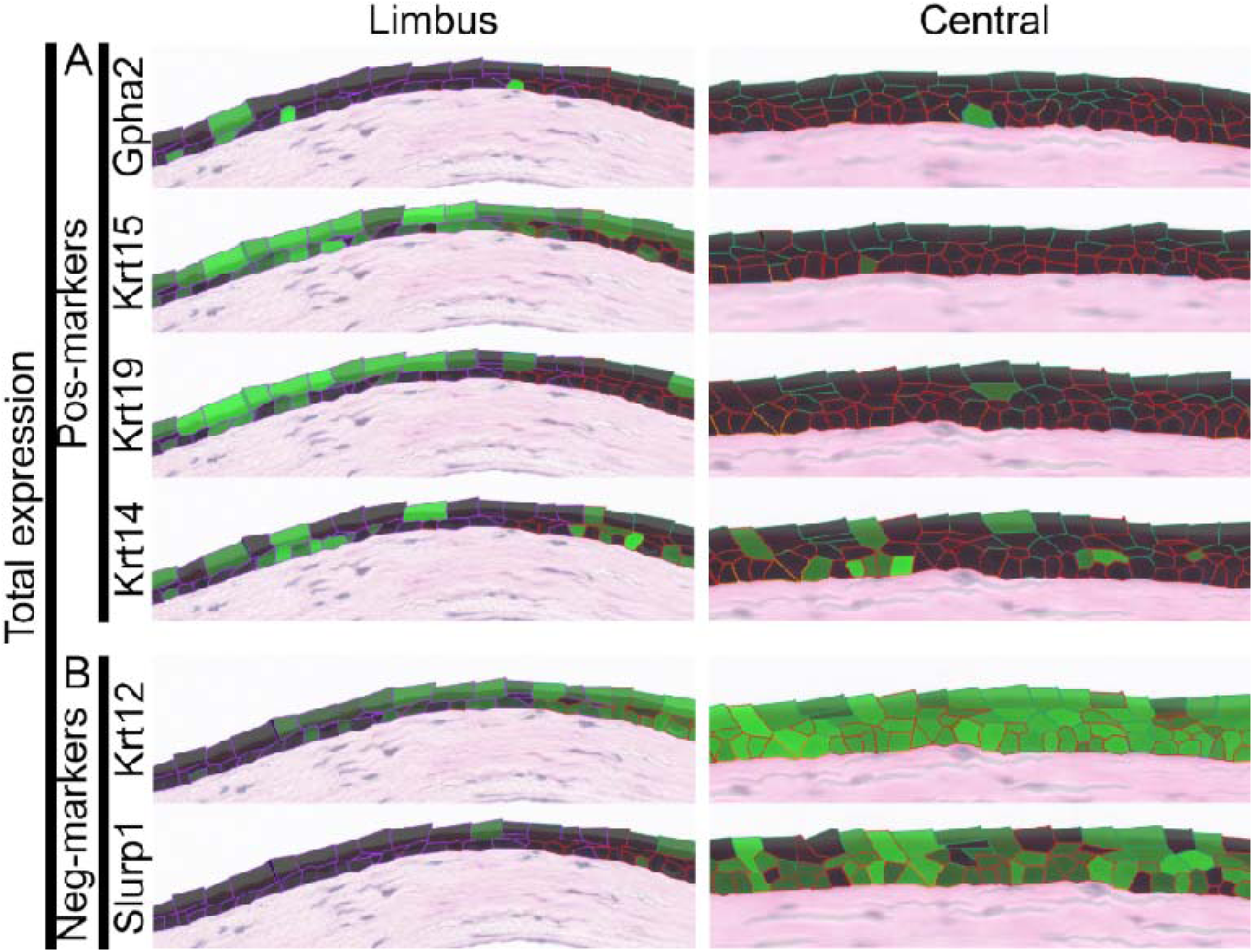
Spatial gene expression maps of LSC-specific positive and negative markers across the epithelium with all epithelial cells shown. Green intensity signifies per-gene normalized expression level within each cell, with maximum intensity mapped to the maximum mean cluster-wise gene expression, and minimum intensity mapped to the minimum mean expression.

**Figure S2.**
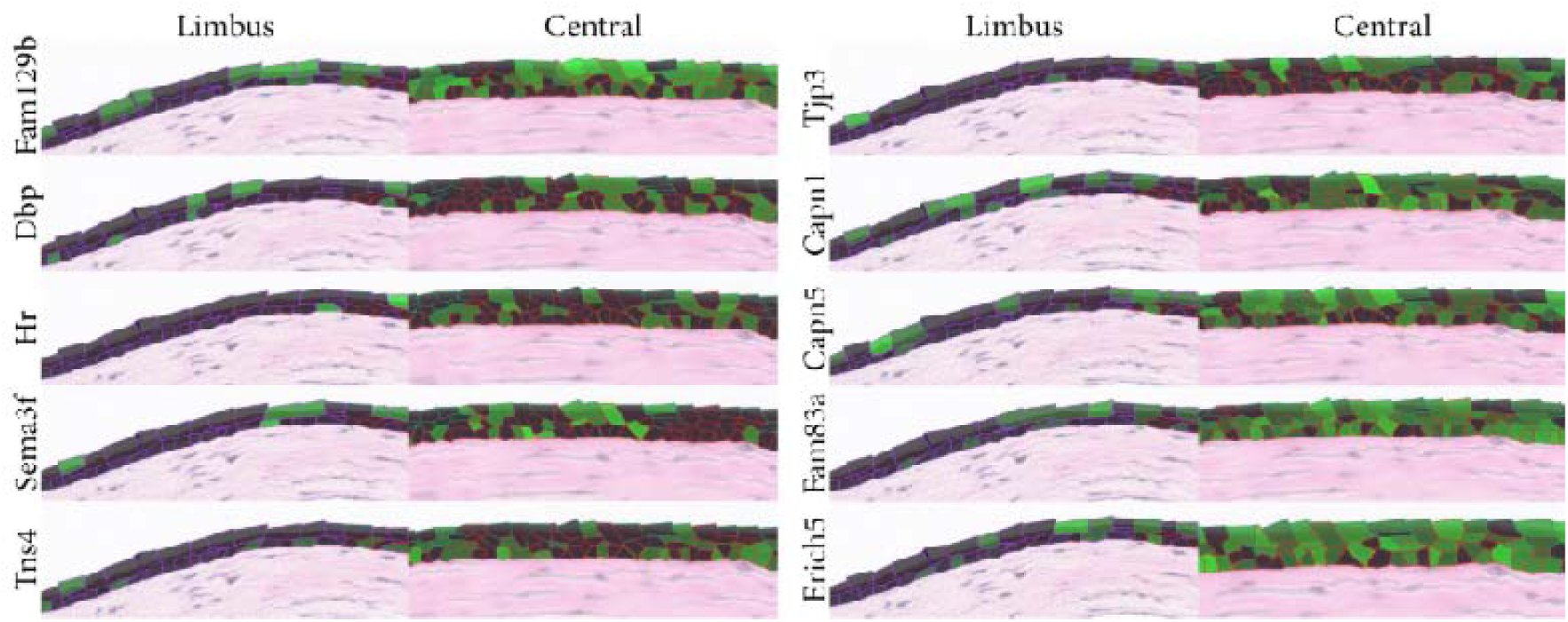
Spatial gene expression maps of ST-enhanced markers across the epithelium. Green intensity signifies per-gene normalized expression level for each cell.

**Figure S3.**
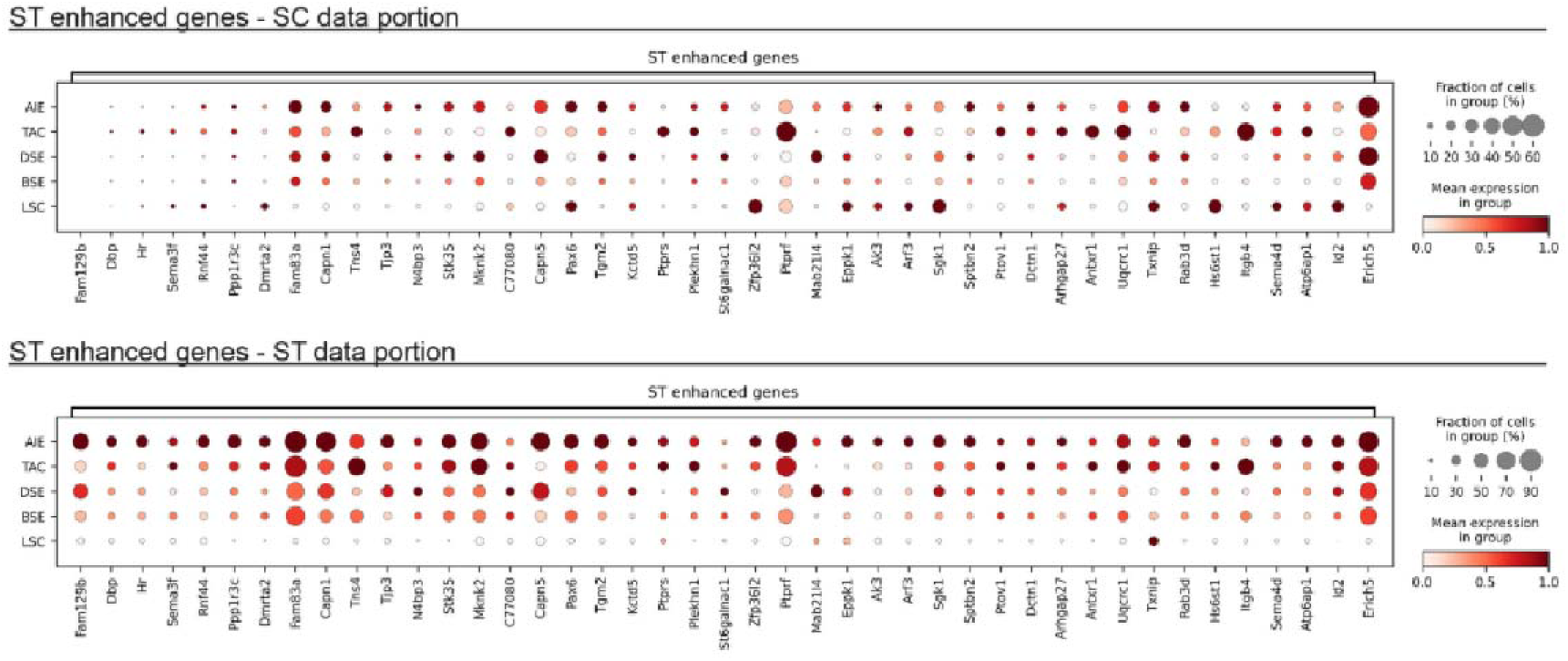
Gene expression of the integration clusters. **a)** Gene expression profile of ST enhanced genes in the SC dataset (top panel) and the ST dataset (bottom panel). Expression values are normalized per gene between 0 (minimum average cluster expression) and 1 (maximum average cluster expression).

**Figure S4.**
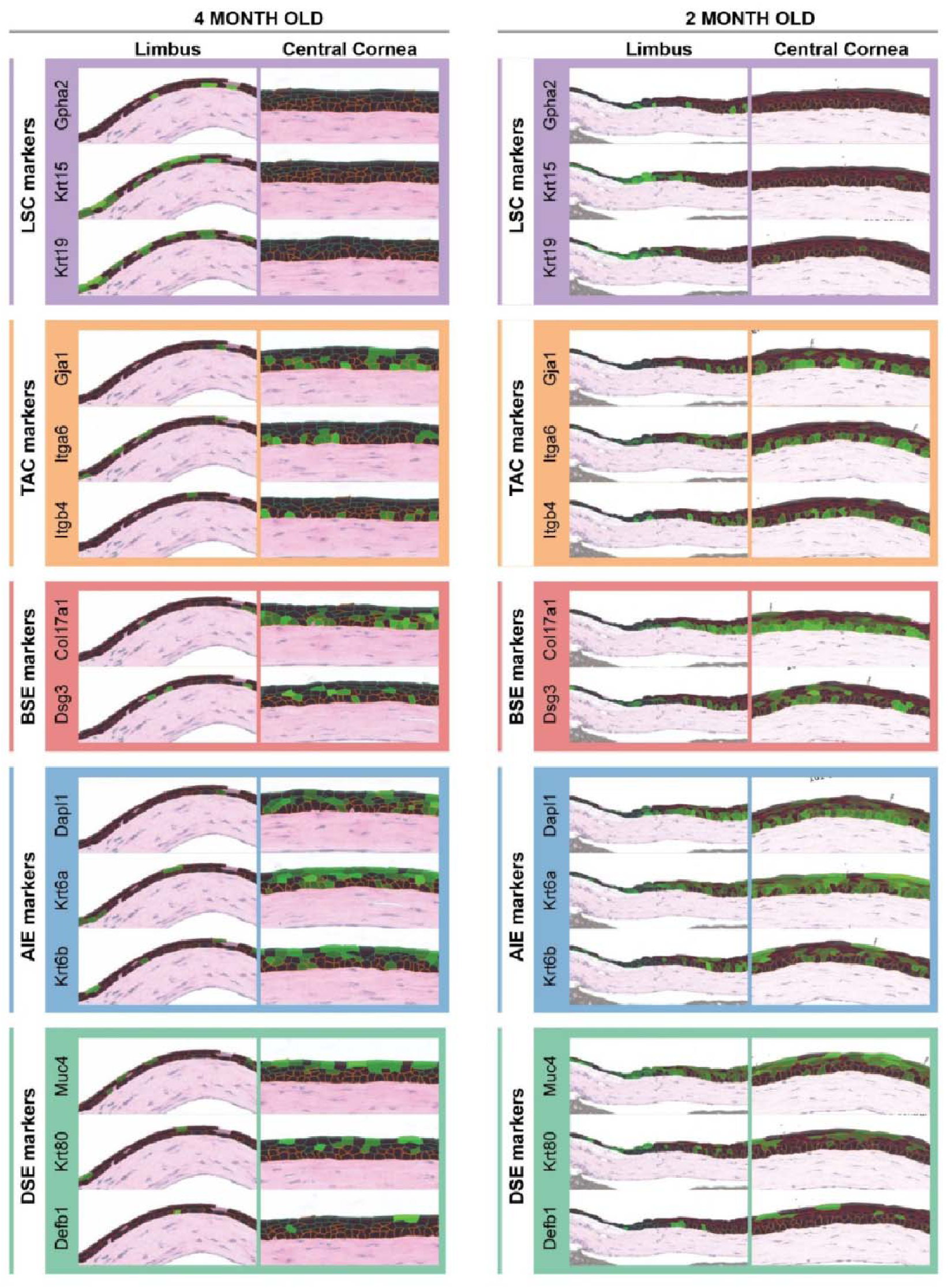
Spatial gene expression maps of integrated cluster-specific markers applied to an independent 4-month-old WT mouse cornea (left column) and younger WT cornea (2 months, right column). Green intensity indicates per-gene normalized expression level within each cell, with the highest intensity mapped to the highest mean cluster-wise gene expression, and minimum intensity mapped to the minimum mean expression.

**Figure S5.**
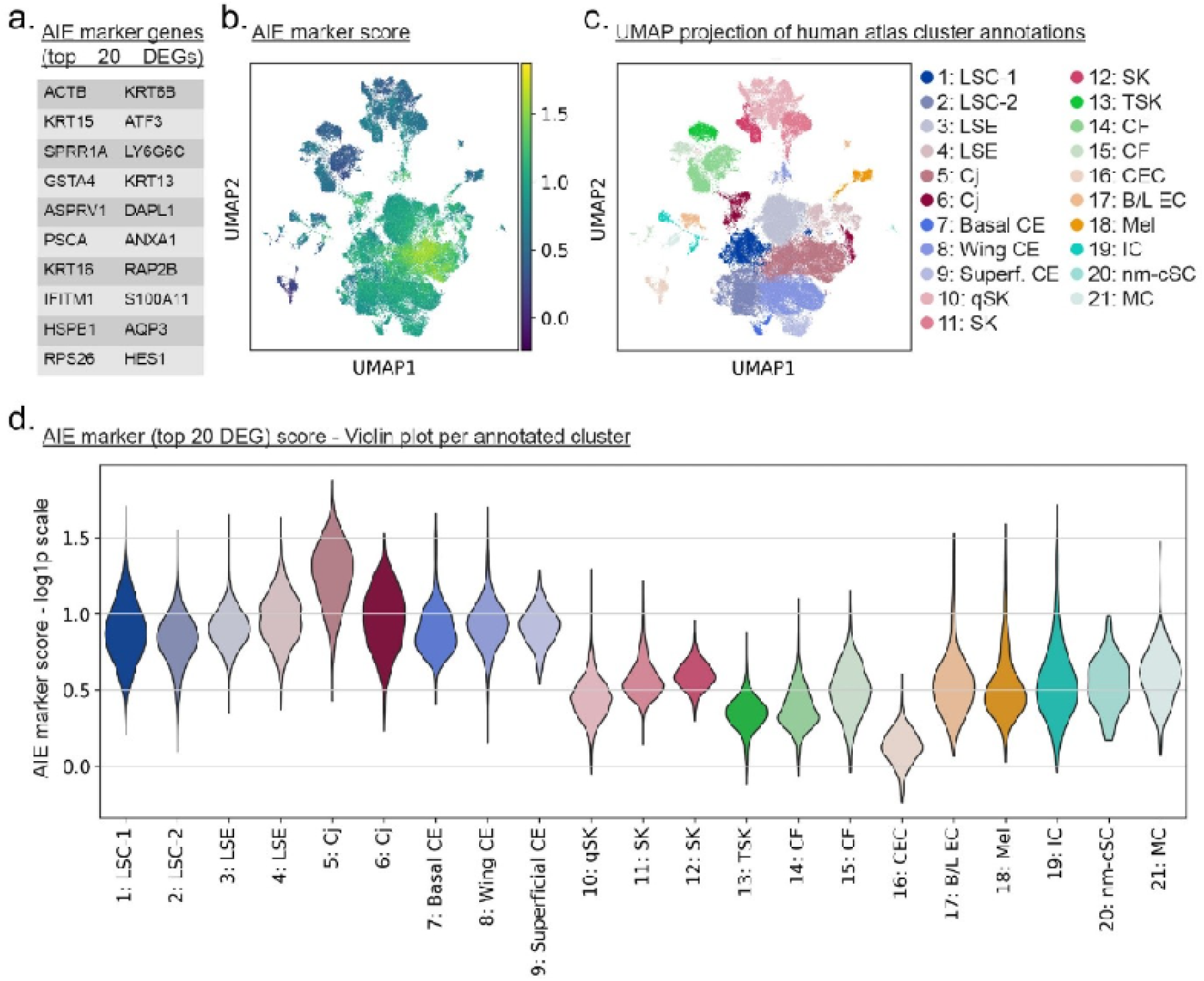
Interrogation of the AIE marker gene set in published human cornea SC datasets. **a)** Top 20 DEG defining the AIE cluster. **b)** UMAP projection of a human cornea scRNA-seq meta-dataset (atlas), with color scale based on similarity score for the top 20 DEGs defining the AIE cluster in our mouse dataset. **c)** UMAP projection of the published cluster annotation on the same UMAP coordinates for cross-correlation. **d)** Violin plot of the calculated AIE marker score of the human dataset, scoring performed with the top 20 DEGs that defined the AIE cluster in our data.

