## Supplementary materials for "Decoding murine corneal epithelial specification and homeostasis by single-cell spatial transcriptomics with scRNA-seq enrichment"

**Supplementary material consists of:**

**Supplementary Table S1**

**Supplementary Table S2**

**Supplementary Table S3**

**Supplementary Table S4**

**Supplementary Figure S1**

**Supplementary Figure S2**

**Supplementary Figure S3**

**Supplementary Figure S4**

**Supplementary Figure S5**

**Table S1. Cluster cell composition after integration.**

| Cluster | ST portion | | | SC portion | | | Enrichment | |
|---|---|---|---|---|---|---|---|---|
| | # of cells | % of integrated ST cells | Average raw counts per cell | # of cells | % of integrated SC cells | Average raw counts per cell | Cluster size increase | Gene count increase |
| **AIE** | 58 | 4.8% | 941.1 | 2904 | 20.6% | 4603.1 | 4907% | 489% |
| **TAC** | 62 | 5.2% | 799.8 | 3935 | 27 .9% | 7308.8 | 6247% | 914% |
| **DSE** | 355 | 29.6% | 613.6 | 3192 | 22.7% | 3465.8 | 799% | 565% |
| **BSE** | 605 | 50.5% | 569.9 | 3664 | 26% | 2916.1 | 506% | 512% |
| **LSC** | 118 | 9.8% | 296.2 | 394 | 2.8% | 4951.5 | 234% | 1672% |

### Table S2. Full list of biological processes enriched across integrated clusters.

| GO ID | Term | Count | P-Value | Fold Enrichment | FDR |
|---|---|---|---|---|---|
| | **LSCs** | | | | |
| **GO:0008285** | negative regulation of cell population proliferation | 14 | 9.09E-08 | 7.13 | 6.41E-05 |
| **GO:0006457** | protein folding | 9 | 2.58E-07 | 14.10 | 9.69E-05 |
| **GO:0015078** | proton transmembrane transport | 9 | 2.75E-07 | 13.99 | 9.69E-05 |
| **GO:0030335** | positive regulation of cell migration | 11 | 9.09E-07 | 8.30 | 0.000257 |
| **GO:0042026** | protein refolding | 5 | 3.14E-06 | 48.12 | 0.000738 |
| **GO:0060261** | positive regulation of transcription by RNA polymerase II | 21 | 5.09E-06 | 3.31 | 0.00088 |
| **GO:0043066** | negative regulation of apoptotic process | 15 | 5.61E-06 | 4.55 | 0.00088 |
| **GO:0009410** | response to xenobiotic stimulus | 11 | 4.95E-06 | 6.86 | 0.00088 |
| **GO:0008380** | RNA splicing | 9 | 7.23E-06 | 9.05 | 0.00102 |
| **GO:0050821** | protein stabilization | 9 | 1.64E-05 | 8.08 | 0.00211 |
| **GO:0051085** | chaperone cofactor-dependent protein refolding | 5 | 2.31E-05 | 29.72 | 0.00272 |
| **GO:0043065** | positive regulation of apoptotic process | 11 | 2.98E-05 | 5.57 | 0.00301 |
| **GO:0051726** | regulation of cell cycle | 9 | 0.000032 | 7.36 | 0.00301 |
| **GO:0045740** | positive regulation of DNA replication | 5 | 2.92E-05 | 28.07 | 0.00301 |
| **GO:0006397** | mRNA processing | 9 | 4.23E-05 | 7.08 | 0.00373 |
| **GO:0030307** | positive regulation of cell growth | 6 | 0.000199 | 11.23 | 0.0162 |
| **GO:0015990** | electron transport coupled proton transport | 3 | 0.000237 | 121.27 | 0.0176 |
| **GO:0030182** | neuron differentiation | 8 | 0.000265 | 6.39 | 0.0181 |
| **GO:0042776** | proton motive force-driven mitochondrial ATP synthesis | 5 | 0.000283 | 15.79 | 0.0181 |
| **GO:0045943** | positive regulation of transcription by RNA polymerase I | 4 | 0.000278 | 31.10 | 0.0181 |
| | **TAC** | | | | |
| **GO:0002181** | cytoplasmic translation | 25 | 3.97E-33 | 45.94 | 5.39E-30 |
| **GO:0140242** | translation at postsynapse | 14 | 7.32E-18 | 43.53 | 3.31E-15 |
| **GO:0140236** | translation at presynapse | 13 | 3.24E-16 | 41.06 | 1.1E-13 |
| **GO:0090136** | cell adhesion | 14 | 2.03E-06 | 5.41 | 0.00055 |
| **GO:0010628** | positive regulation of gene expression | 14 | 6.94E-06 | 4.83 | 0.00135 |
| **GO:0030335** | positive regulation of cell migration | 9 | 5.69E-05 | 6.79 | 0.0068 |
| **GO:0045471** | response to ethanol | 7 | 6.67E-05 | 10.18 | 0.00696 |
| **GO:0000028** | ribosomal small subunit assembly | 4 | 7.51E-05 | 47.56 | 0.00728 |
| **GO:0042274** | ribosomal small subunit biogenesis | 6 | 0.000084 | 13.47 | 0.0076 |
| **GO:0045471** | response to ethanol | 7 | 6.67E-05 | 10.18 | 0.00696 |
| **GO:0000028** | ribosomal small subunit assembly | 4 | 7.51E-05 | 47.56 | 0.00728 |
| **GO:0042274** | ribosomal small subunit biogenesis | 6 | 0.000084 | 13.47 | 0.0076 |
| **GO:0010629** | negative regulation of gene expression | 10 | 0.000163 | 5.08 | 0.0139 |
| **GO:0031581** | hemidesmosome assembly | 3 | 0.000355 | 101.06 | 0.0283 |
| **GO:0045785** | positive regulation of cell-cell adhesion | 4 | 0.000428 | 26.95 | 0.0323 |
| **GO:0045944** | positive regulation of transcription by RNA polymerase II | 17 | 0.000597 | 2.68 | 0.0427 |
| **GO:0001666** | response to hypoxia | 7 | 0.000735 | 6.52 | 0.0499 |
| | **BSE** | | | | |
| **GO:0006338** | chromatin remodeling | 14 | 1.84E-05 | 4.40 | 0.015 |
| | **AIE** | | | | |
| **GO:0045109** | intermediate filament organization | 8 | 5.31E-08 | 23.26 | 5.24E-05 |
| **GO:0031424** | keratinization | 6 | 3.56E-05 | 16.12 | 0.0176 |
| **GO:0030336** | negative regulation of cell migration | 7 | 8.03E-05 | 9.84 | 0.0265 |
| **GO:0030855** | epithelial cell differentiation | 6 | 0.000165 | 11.68 | 0.0407 |
| | **DSE** | | | | |
| **GO:0031424** | keratinization | 8 | 9.22E-08 | 21.49 | 7.25E-05 |
| **GO:0045109** | intermediate filament organization | 7 | 1.24E-06 | 20.35 | 0.00486 |

**Table S3. ST-enhanced genes.** The full list of 43 ST-enhanced genes, based on minimum total log-normalized expression level of 0.5 and > 3-fold ST/SC expression ratio.

| No | Gene | ST expression | SC expression | ST/SC ratio | No | Gene | ST expression | SC expression | ST/SC ratio |
|---|---|---|---|---|---|---|---|---|---|
| **1** | Fam129b | 1.28 | Not detectable | N/A | **22** | St6galnac1 | 0.53 | 0.13 | 3.92 |
| **2** | Dpb | 0.58 | 0.02 | 23.58 | **23** | Zfp362 | 0.61 | 0.16 | 3.92 |
| **3** | Hnr | 0.66 | 0.04 | 14.87 | **24** | Ptprf | 1.65 | 0.42 | 3.90 |
| **4** | Sema4f | 0.61 | 0.05 | 12.00 | **25** | Mab21l4 | 0.61 | 0.16 | 3.83 |
| **5** | Rnf44 | 0.73 | 0.08 | 9.45 | **26** | Eppl1 | 0.65 | 0.17 | 3.77 |
| **6** | Ppp1r3c | 0.64 | 0.08 | 7.83 | **27** | Ak3 | 0.55 | 0.15 | 3.70 |
| **7** | Dmrt2a | 0.54 | 0.07 | 7.56 | **28** | Arf3 | 0.54 | 0.15 | 3.64 |
| **8** | Fam83a | 2.29 | 0.31 | 7.51 | **29** | Sgk1 | 0.81 | 0.23 | 3.59 |
| **9** | Capn1 | 1.53 | 0.22 | 6.84 | **30** | Sptbn2 | 0.62 | 0.18 | 3.55 |
| **10** | Tns4 | 1.15 | 0.18 | 6.33 | **31** | Pvt1 | 0.52 | 0.15 | 3.54 |
| **11** | Tip3 | 0.90 | 0.15 | 6.07 | **32** | Dctn1 | 0.52 | 0.15 | 3.47 |
| **12** | N4bp3 | 0.62 | 0.10 | 6.07 | **33** | Arhgap2 | 0.55 | 0.16 | 3.46 |
| **13** | Stk36 | 0.99 | 0.18 | 5.36 | **34** | Antxr1 | 0.55 | 0.16 | 3.46 |
| **14** | Mknk2 | 1.33 | 0.26 | 5.13 | **35** | Uqcrc1 | 1.00 | 0.29 | 3.43 |
| **15** | C77080 | 0.69 | 0.15 | 4.70 | **36** | Txnip | 0.75 | 0.22 | 3.39 |
| **16** | Caps6 | 1.44 | 0.31 | 4.61 | **37** | Rab3d | 0.66 | 0.20 | 3.32 |
| **17** | Pax6 | 1.03 | 0.24 | 4.25 | **38** | Hs6st4 | 0.53 | 0.16 | 3.31 |
| **18** | Tgm2 | 0.86 | 0.20 | 4.23 | **39** | Itgb4 | 0.84 | 0.25 | 3.31 |
| **19** | Kctd5 | 0.51 | 0.12 | 4.21 | **40** | Sema4d | 0.56 | 0.17 | 3.27 |
| **20** | Ptprs | 0.51 | 0.13 | 4.01 | **41** | Atp6ap1 | 0.55 | 0.17 | 3.19 |
| **21** | Plekhh1 | 0.56 | 0.14 | 3.93 | **42** | Id2 | 0.68 | 0.22 | 3.09 |
| | | | | | **43** | Erich5 | 2.08 | 0.67 | 3.08 |

**Table S4. ST-enhanced biological processes.** GO term enrichment of spatially emphasized biological processes, based on unadjusted p-value < 0.05 and ST-enhanced gene list and minimum count of three genes.

| GO ID | Term | Genes | P-value | Fold Enrichment | FDR |
|---|---|---|---|---|---|
| **GO:0001755** | neural crest cell migration | Pax6; Sema3f; Sema4d | 0.00272 | 37.95 | 0.61 |
| **GO:0010975** | regulation of neuron projection development | Pax6; Ptprf; Ptprs | 0.00272 | 37.95 | 0.61 |
| **GO:0007399** | nervous system development | N4bp3; Pax6; Ptprf; Ptprs; Sema4d | 0.00704 | 6.38 | 0.983 |
| **GO:0009611** | response to wounding | Itgb4; Pax6; Zfp36l2 | 0.0123 | 17.41 | 0.983 |
| **GO:0043065** | positive regulation of apoptotic process | Plekhh1; Ptprf; Txnip; Tgm2 | 0.0219 | 6.52 | 1 |
| **GO:0007411** | axon guidance | Pax6; Sema3f; Sema4d | 0.0368 | 9.68 | 1 |

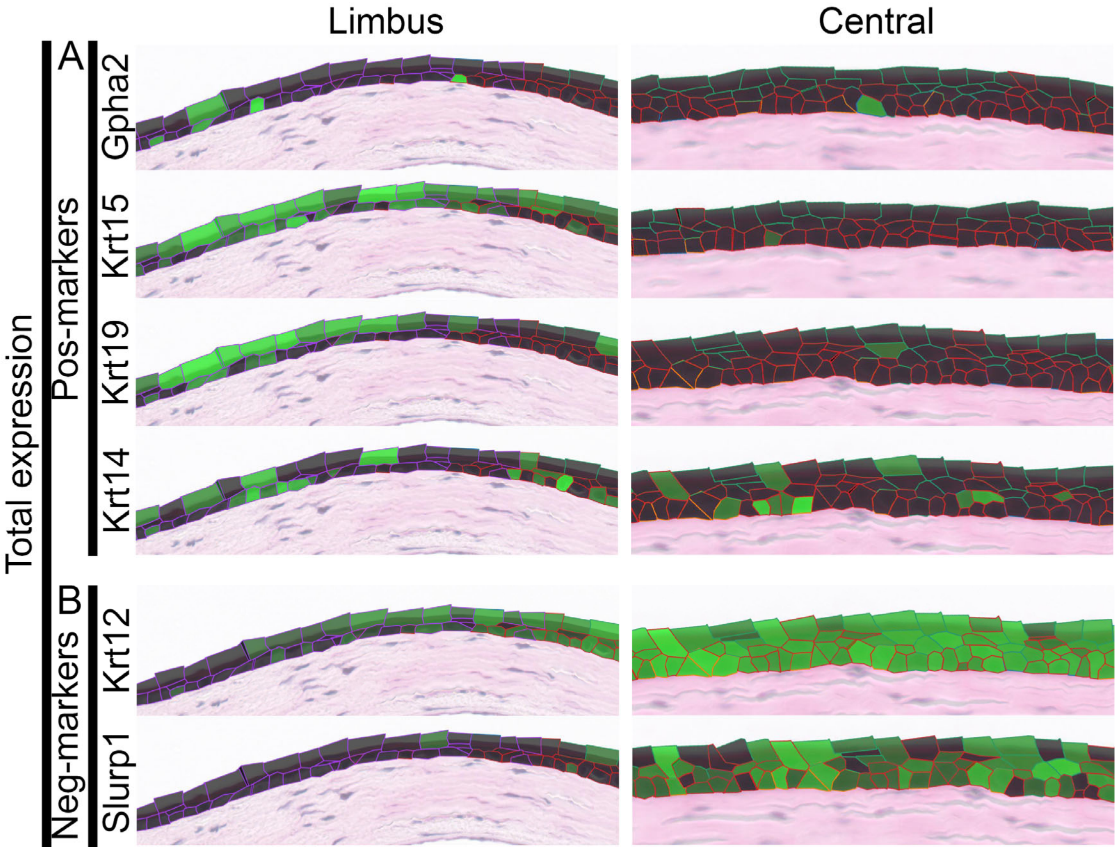


**Figure S1.** Spatial gene expression maps of LSC-specific positive and negative markers across the epithelium with all epithelial cells shown. Green intensity signifies per-gene normalized expression level within each cell, with maximum intensity mapped to the maximum mean cluster-wise gene expression, and minimum intensity mapped to the minimum mean expression.

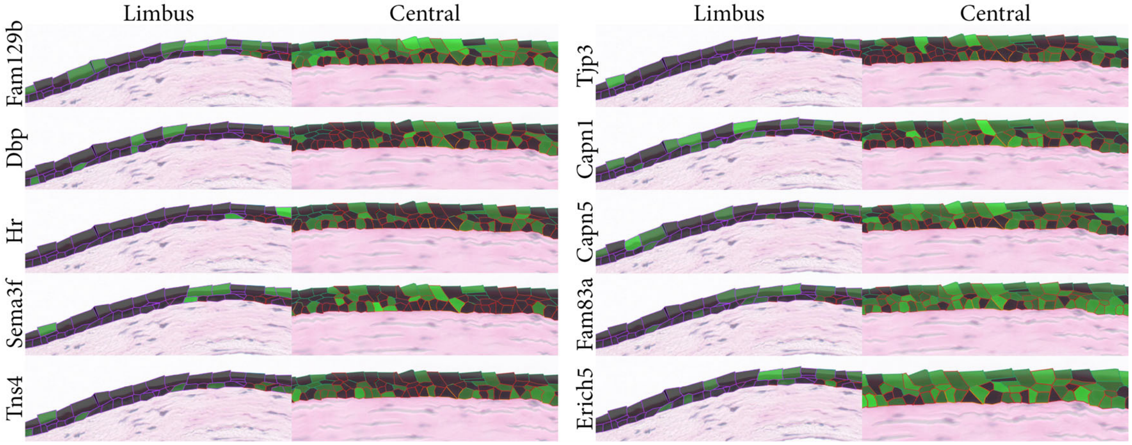


**Figure S2.** Spatial gene expression maps of ST-enhanced markers across the epithelium. Green intensity signifies per-gene normalized expression level for each cell.

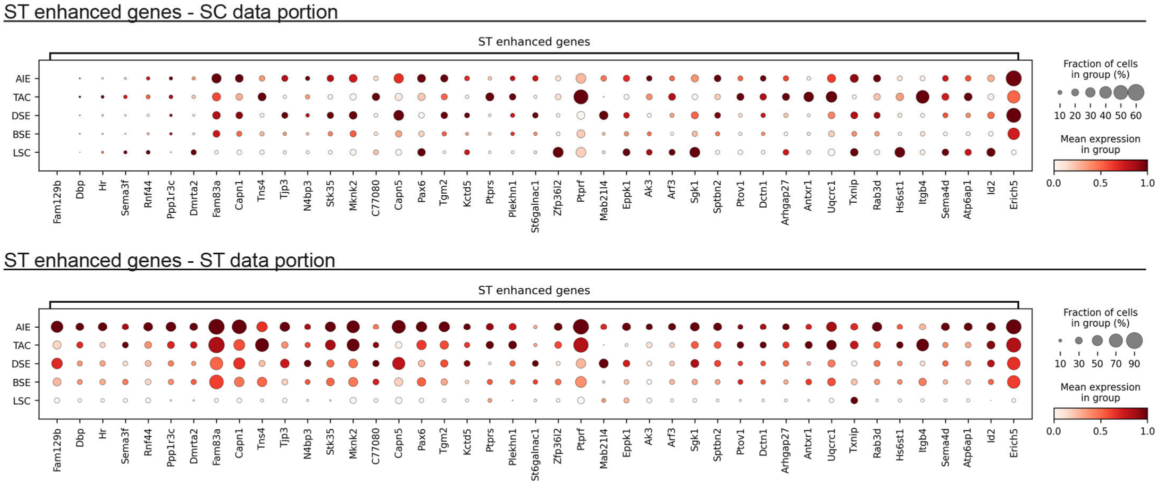


**Figure S3. Gene expression of the integration clusters. a)** Gene expression profile of ST enhanced genes in the SC dataset (top panel) and the ST dataset (bottom panel). Expression values are normalized per gene between 0 (minimum average cluster expression) and 1 (maximum average cluster expression).

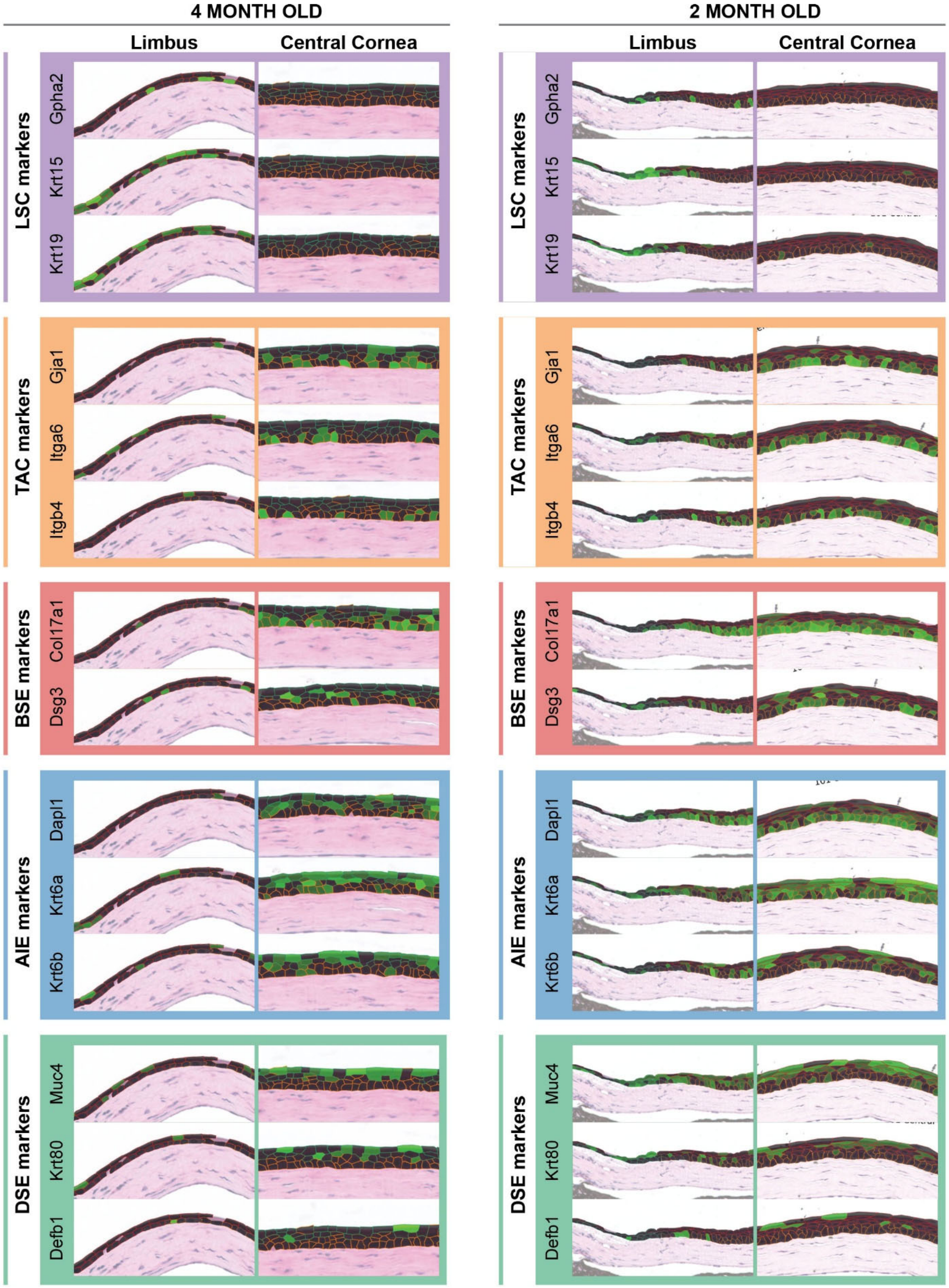


**Figure S4.** Spatial gene expression maps of integrated cluster-specific markers applied to an independent 4-month-old WT mouse cornea (left column) and younger WT cornea (2 months, right column). Green intensity indicates per-gene normalized expression level within each cell, with the highest intensity mapped to the highest mean cluster-wise gene expression, and minimum intensity mapped to the minimum mean expression.

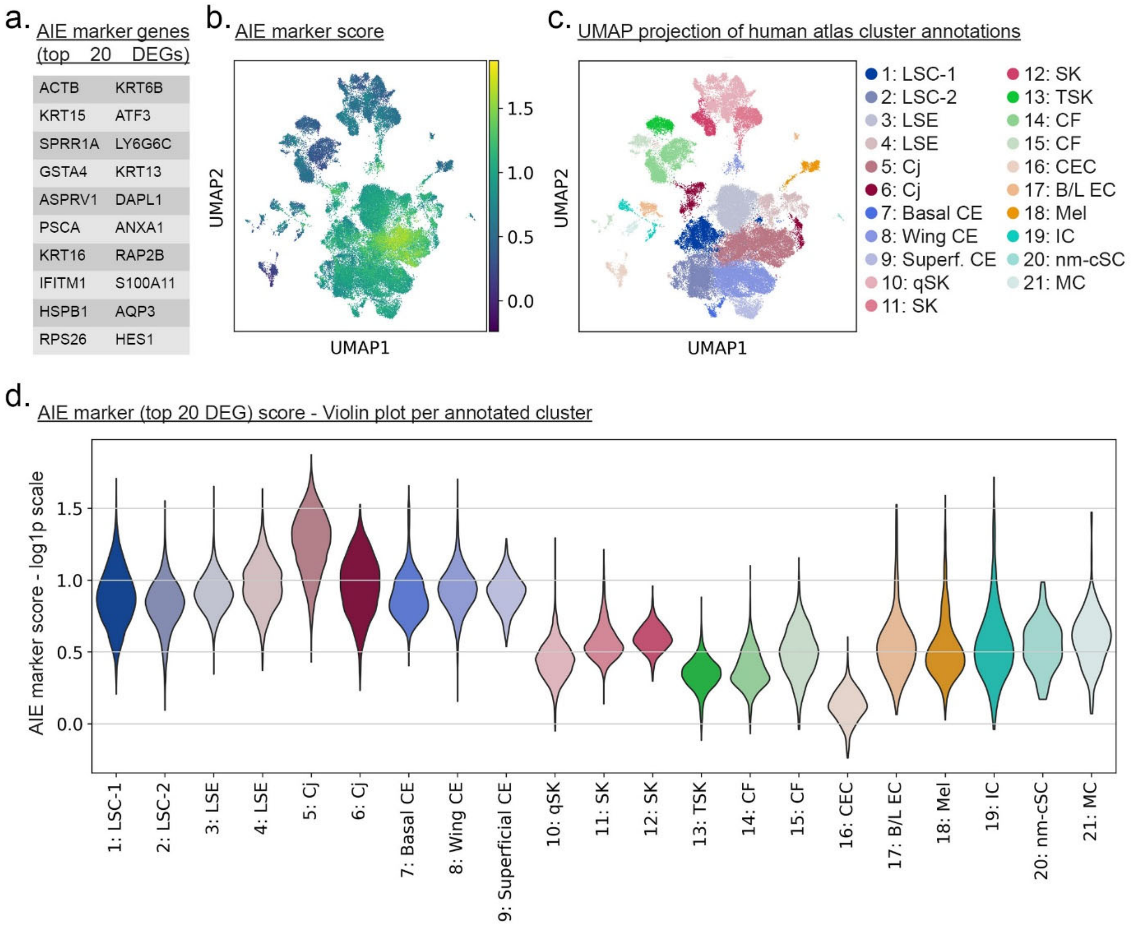


**Figure S5. Interrogation of the AIE marker gene set in published human cornea SC datasets. a)** Top 20 DEG defining the AIE cluster. **b)** UMAP projection of a human cornea scRNA-seq meta-dataset (atlas), with color scale based on similarity score for the top 20 DEGs defining the AIE cluster in our mouse dataset. **c)** UMAP projection of the published cluster annotation on the same UMAP coordinates for cross-correlation. **d)** Violin plot of the calculated AIE marker score of the human dataset, scoring performed with the top 20 DEGs that defined the AIE cluster in our data.
